# Design of colipids with controlled lateral distance of head group moieties for targeting strategies in the field of lipid-based nanoparticles

**DOI:** 10.64898/2026.11.09.765079

**Authors:** Vincent Kampik, Sebastian Ammenwerth, Matthias Elgeti, Christian Wölk

**Affiliations:** Pharmaceutical Technology, Institute for Pharmacy, Medical Faculty, Leipzig University, Eilenburger Straße 15a, 04317 Leipzig, Germany; Institute for Drug Discovery and Institute for Medical Physics and Biophysics, University of Leipzig Medical School, Leipzig, Germany; Integrative Center for Bioinformatics, University of Leipzig, Leipzig, Germany

## Abstract

Lipid nanoparticles and liposomes are nanoparticle drug formulations which are firmly established in clinics. In order to expand their application and improve efficacy, surface modifications can be a key strategy, especially for cell specific targeting. Targeting can be achieved by integrating ligand functionalized lipids or lipidoids on the nanoparticle surface. In this case, not solely the targeting ligand itself, but also the structure of the whole targeting lipid/lipidoid can affect the interaction of the targeting ligand with the binding motive on cells. Especially, distal spacer structures between lipid anchor and ligand unit, often composed of polyethylene glycol chains, provide a certain distance of the ligand from the nanoparticle surface. A concept which is currently neglected is the control of the lateral distance between ligand structures on the nanoparticle. Herein we investigated the possibility to design lipidoids with a lateral spacer composed of polyethylene glycol units. We synthesized and characterized different lipidoids which provide a primary amine in the head group for possible ligand coupling. In our model lipidoids the head groups were functionalized with a paramagnetic probe to apply electron paramagnetic resonance (EPR) methods for the characterization of the lipidoid interaction with model nanoparticles, in our case liposomes. Continuous wave EPR was used to study the efficiency of the novel lipidoids to integrate into lipid bilayers of model liposomes, which was shown to depend on lipidoid structure and liposome composition. Furthermore, using double electron-electron resonance (DEER), we observed lipidoid conformational changes occurring upon integration into liposomes.

**Graphical Abstract:** 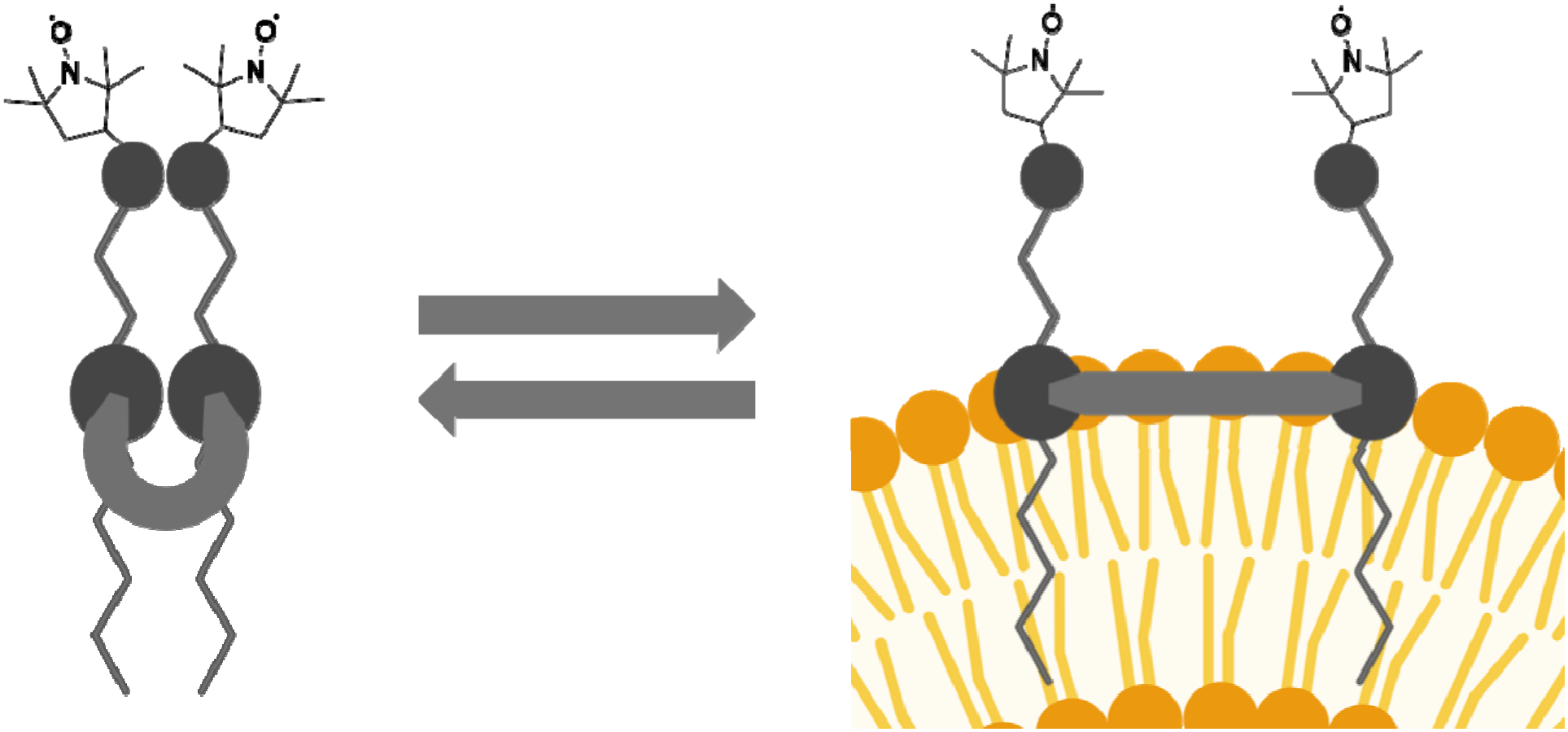

## 1. Introduction

Lipid-based nanocarriers are an established delivery system for therapeutic molecules, with the first FDA approved medicine being doxorubicin encapsulated in liposomes.^1^ Another type of lipid-based nanocarriers are nucleic acid encapsulating lipid nanoparticles (LNPs) used for approved drugs like Onpatro^TM^ or Comirnati^TM^. But still, broad and unspecific biodistribution resulting in off target effects are a challenge for the development of LNP or liposome formulations.^2^ One strategy of overcoming the problem of non-specific distribution is the use of active targeting. Here, targeting ligands on the lipid-based nanoparticle surface are utilized to promote internalization by specific cell types, and therefore improving their specificity.^3^ Conjugates of the ligand with lipids are used to present the ligand on the nanoparticle surface. These targeting lipids usually have three subunits. The lipid part, the ligand and a spacer (usually polyethylene glycol (PEG)) connecting the two first mentioned subunits.^4^ There are different options for the targeting ligand, e.g., polysaccharides, peptides, aptamers or small molecules, but the usual structure of the targeting lipids only allows one ligand per lipid.^5^

In this work we present two model targeting lipidoids as a proof-of-concept for a new lipid platform, that allows the use of two targeting ligands per lipid, creating a controlled distance between those binding motifs. This concept is shown in Figure 1. The strategy can hypothetically improve efficiency of the LNP formulation, since the pairing of two binding motifs in a certain distance can improve cell uptake.^6^ Also a higher specificity of binding is a possible effect of such lateral distance control, as some receptor – co-receptor pairings are only present in certain cell types, such as the CD4/CCR5 pairing for T-helper cells.^7^ Improved efficacy and specificity would allow a dose reduction of LNP formulations, which may lead to less side effects.^8^

**Figure 1:**
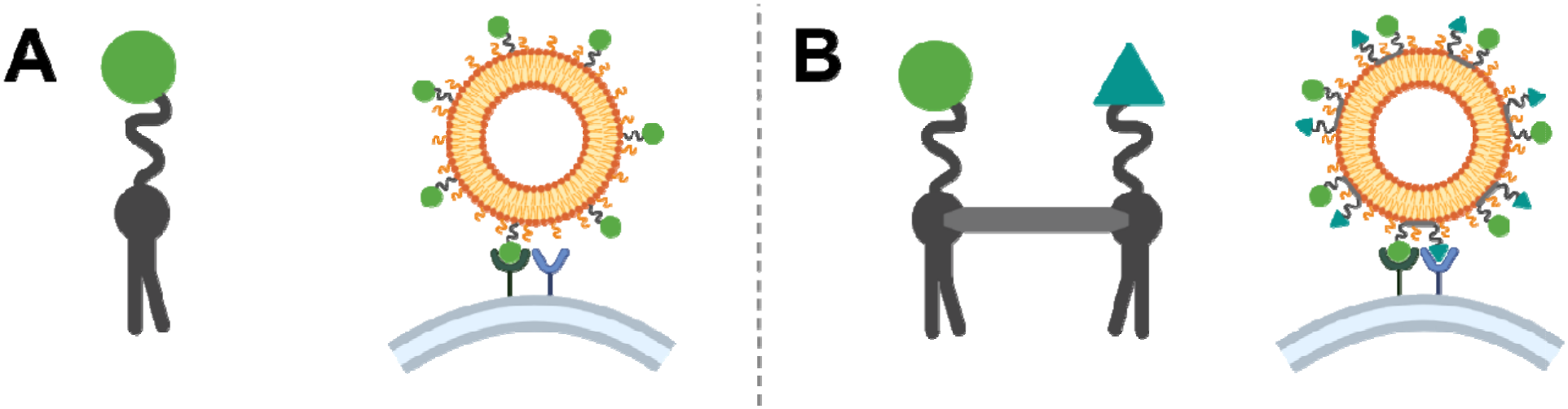
comparison of (A) monovalent and (B) bivalent targeting lipids/lipidoids and their respective binding to targets on cell surfaces

A key challenge in the use of targeting lipids is the formulation process, since these lipids can affect important characteristics of the lipid-based nanoparticles like size or zeta potential.^9^ Another concern is the localization of the targeting lipid, more precisely, the exclusive presence on the nanoparticle surface, a requirement for binding to the target cell.^4^ The most common method to address these challenges is the “post-insertion” method to decorate lipid-based nanoparticles with target ligands. In this strategy the nanoparticle gets prepared without the targeting moiety and the targeting lipid is added to the formulation as micellar or molecular solution followed by an insertion into the lipid-based nanoparticle surface driven by partition/adsorption effects.^10^ Most post-insertion protocols use an incubation time of about 60 min, making the method very quick.^11–13^ It also allows a lot of flexibility, since different targeting lipids can be inserted in the same pre-existing formulation.^14^ With optimized and strictly controlled conditions, it is also possible to achieve a functionalization of the LNP or liposome formulation exclusively on the outer surface.^15^ However, using the post-insertion method results in the question which amount of the targeting lipid is present in the final lipid-based nanoparticle formulation, since the method relies on the self-insertion of the targeting lipid. Multiple methods have been described in literature to address this issue, like weighing the dried residue or size exclusion chromatography with fluorescence labeled liposomes and post-inserted lipid.^10,16^ But also spectroscopic techniques can help to understand the mechanism of post-insertion.

In this work we apply the method of post-insertion on the functionalization of liposomes with a new species of bivalent targeting lipidoids characterized by a lateral spacer to control the distance between the two head groups. We quantify the amount of post-inserted targeting lipidoid using continuous wave electron paramagnetic resonance (CW-EPR). Additionally, we use double electron-electron resonance (DEER) to check for the ability of the polyethylene glycol based lateral spacer to control the distance between the two intramolecular head groups of the lipidoid.

## 2. Results and discussion

### 2.1. Synthesis model lipidoids

The synthesis of the two model-lipidoids (see structures Figure 2) followed a synthetic strategy developed by Wölk et al. for the preparation of peptide-mimic lysine-based amino-functionalized lipids, which bases on distinct building blocks.^17^ Two lysine-based lipidoid units were conjugated via a PEG-spacer, resulting in a bivalent lipidoid bearing two free amino groups in the lipidoid head, serving as attachment sites for potential targeting moieties. In the case of this study, PROXYL was selected as a model head group instead of a targeting moiety allowing the final lipidoids to be analyzed using EPR spectroscopy. Since common targeting lipids in literature use 1,2-distearoyl-*sn*-glycero-3-phosphoethanolamine as lipid part, C16 was selected as length for the alkyl chains.^18–20^ Two synthetic routes were established with two different PEG-spacers: a PEG-diamine (resulting lipid: K16-PEG-N-PROXYL; Figure 2) and a dicarboxylic acid-PEG (resulting lipid: K16-PEG-S-PROXYL; Figure 2). The modular design of the compounds offers a key advantage, as the individual lipidoid components can be synthesized separately and connected via amide bond formation. This enables combinatorial variation of the alkyl chain, spacer and targeting group of the lipidoids, facilitating further structural optimization and modification.

**Figure 2:**
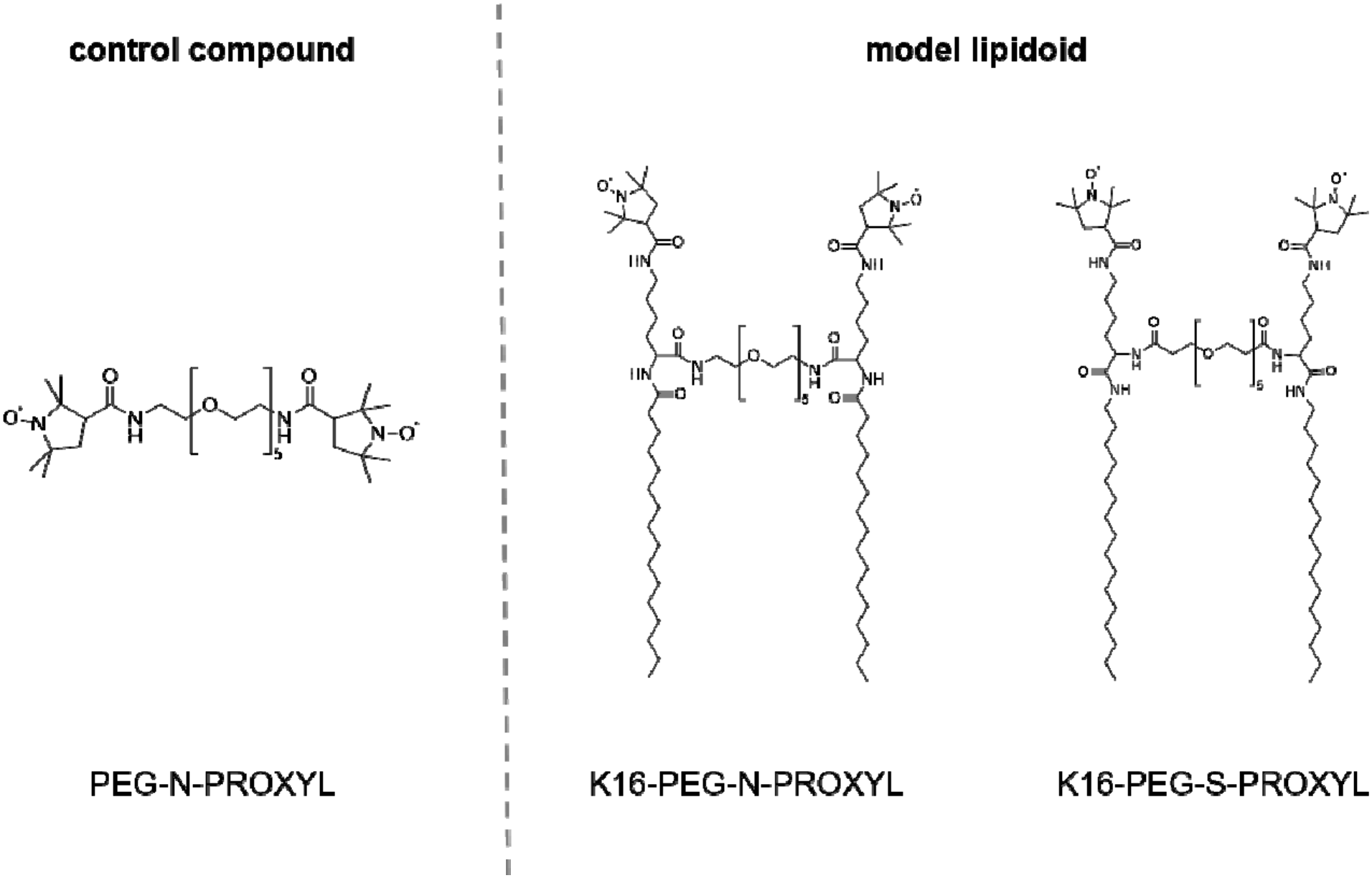
Chemical structure and abbreviations of the two PROXYL labeled model lipidoids and the control compoun

For the synthesis of K16-PEG-N-PROXYL (details Scheme 1), two equivalents of an orthogonal protected L-lysine derivative bearing an amine labile 9-fluorenylmethoxy-carbonyl (Fmoc) protective group in α-position and an acid labile *tert*-butoxycarbonyl (Boc) protective group in ε-position respectively, were coupled to the amino-PEG spacer **1** via amide bond formation yielding in the lysine-derivatized PEG **3**. Fmoc cleavage with piperidine yielded compound **4** with free amino-groups in α-position of the two lysine subunits, the attachment points for coupling of the lipophilic palmitic acid to synthesize compound **5**. Acid induced Boc cleavage resulted in the bivalent amino lipidoid **6**. T make the compound accessible to EPR-measurements, PROXYL-COOH was coupled to the two amino-headgroups yielding the model lipidoid K16-PEG-N-PROXYL.

**Scheme 1:**
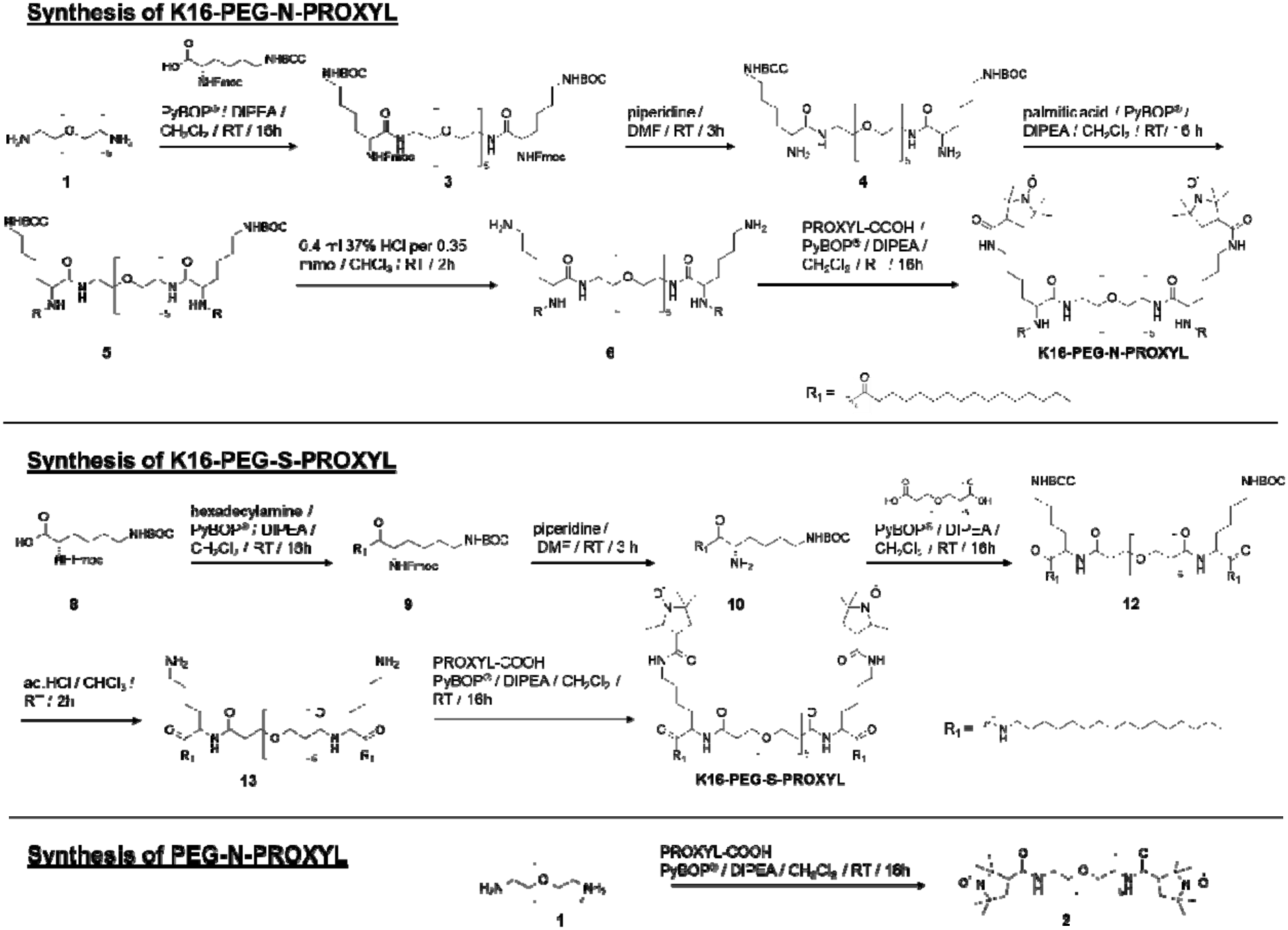
Synthesis of K16-PEG-N-PROXYL, K16-PEG-S-PROXYL, and PEG-N-PROXYL. Abbreviation of reagents and conditions: PyBOP - benzotriazol-1-yloxytripyrrolidinophosphonium hexafluorophosphate; DIPEA -N,N-diisopropylethylamine; RT – room temperature; DMF – dimethylformamide; PROXYL-COOH-3-(carboxy)-2,2,5,5-tetramethyl-1-pyrrolidinyloxy

As starting point for the synthesis of K16-PEG-S-PROXYL the orthogonally protected L-lysine derivative **8** was functionalized with an alkyl residue via amide coupling of hexadecyl amine yielding in the orthogonally protected lysine amide **9** (Scheme 1). The piperidine-catalyzed Fmoc deprotection yielded the lysine amide **10** with the free ε-amine. Two equivalents of the compound **10** were coupled to the PEG-acid spacer 4,7,10,13,16-pentaoxanonadecanedioic acid (compound **11**) to yield the lipidoid precursor **12**. The hydrochloric acid induced Boc-cleavage resulted in the bivalent amino lipidoid **13**. To achieve EPR-activity, PROXYL-COOH was coupled to the two amino-head groups yielding the model lipidoid K16-PEG-S-PROXYL.

To get a reference structure for the EPR spectroscopy that is unable to insert into the liposome surface due to the absence of alkyl chains, a PROXYL labelled version of the diamino-PEG (resulting compound: PEG-N-PROXYL; Figure 2) as a control compound was synthesized (see Scheme 1).

### 2.2. Physico-chemical characterization of model-lipidoids

The model-lipidoids and the control compound were characterized regarding physico-chemical properties to correlate them with the yield of post-insertion, hence, critical micelle concentration (CMC) and the self-aggregating tendencies of targeting lipids are considered key factors for the post-insertion process.^21,22^ To keep comparability between the experiments, we have chosen 10% glycerol as solvent, because it was necessary as a cryo-protector for DEER measurements, the key analytical technique to determine lateral distance control in the lipidoids.

#### 2.2.1. Size measurement in aqueous dispersion of the model lipidoids

To investigate the presence of possible self-associates of the compounds in solution, DLS measurements were performed at 1 mg/mL lipoid dispersions (standard concentration for initial testing in our laboratory), resulting in final concentrations of 1621 µM for PEG-N-PROXYL, 742 µM for K16-PEG-N-PROXYL, and 726 µM K16-PEG-S-PROXYL). Further we have chosen 50 µM lipidoid in aqueous solution as second concentration for the size measurement, a value equal to the used post-insertion conditions. The curves for the volume-weighted size distribution are presented in Figure 3A-C**Error! Reference source not found.**. At the high concentration all three compounds showed particles in the colloidal range (0.5 to 1000 nm). Interestingly at 50 µM, the concentration for the post-insertion, only the solution of the control compound PEG-N-PROXYL consisted of colloidal particles. In contrast, for the two model lipidoids K16-PEG-N-PROXYL and K16-PEG-S-PROXYL, sizes below 1 nm were detected indicating molecular solution. The absence of larger particles could be an advantage for post-insertion, since it is considered to be beneficial to work below the concentration of self-assembly of the targeting lipid.^22^

**Figure 3:**
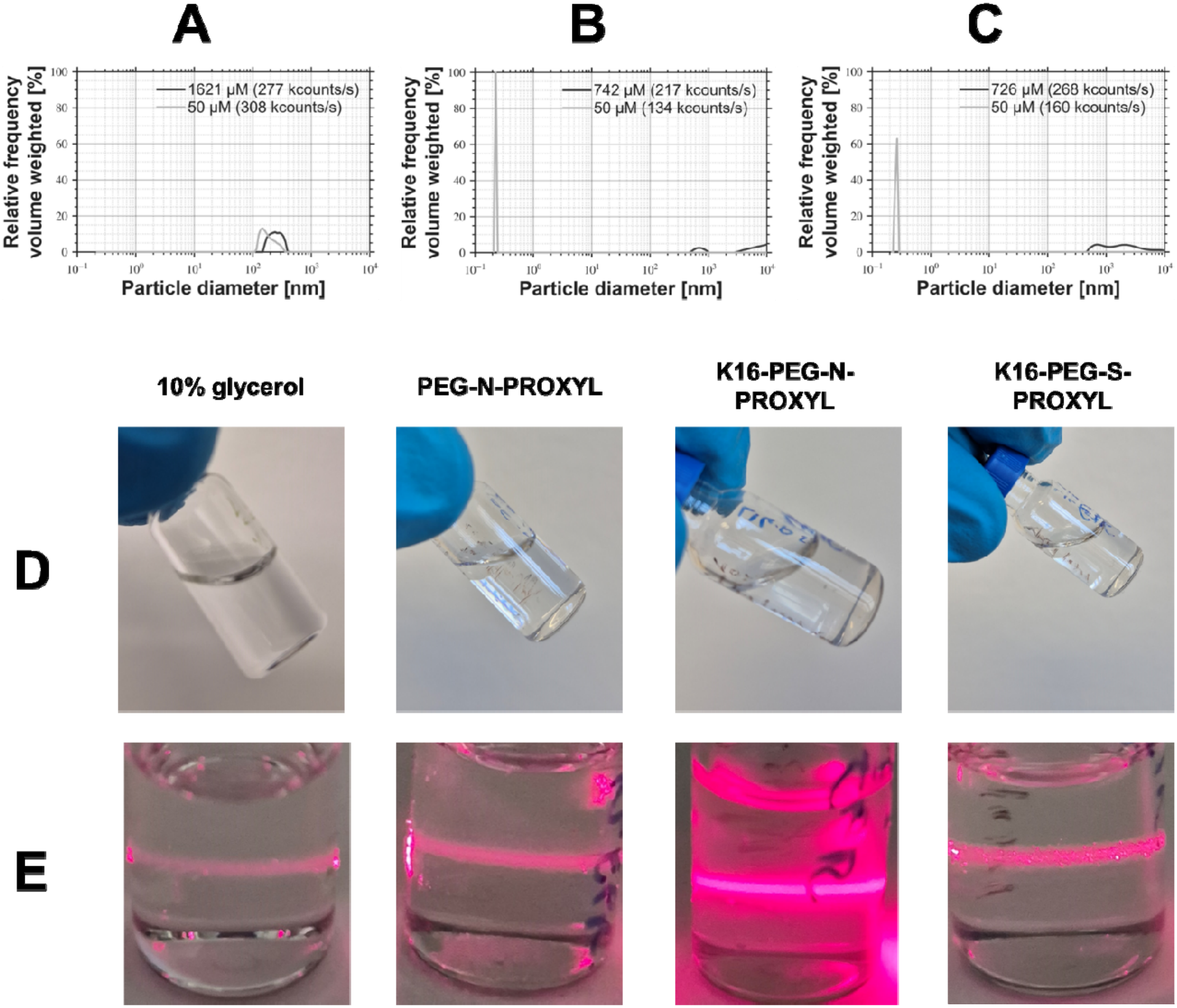
(A-C) volume-weighted size distribution curves at 1 mg/mL (1621 µM for PEG-N-PROXYL; 742 µM for K16-PEG-N-PROXYL; 726 µM for K16-PEG-S-PROXYL) and 50 µM solutions of (A) PEG-N-PROXYL (B) K16-PEG-N-PROXYL (C) K16-PEG-S-PROXYL. All measurements have been performed in 10% glycerol. (D) images of compound solutions in 10% glycerol solution and (E) images of Tyndal effect of the compounds in 10% glycerol solution at the high concentration of each model lipid (1 mg/mL).

For macroscopic evaluation of the colloidal solutions, we additionally checked for the Tyndall effect (TE). The TE arises when light hits a colloidal solution with particles smaller than the wavelength of the incident light, rendering the laser beam visible.^23^ Images of the model lipidoids in solutions (1 mg/mL in 10% glycerol) upon laser illumination (λ = 630 – 680 nm) are shown in Figure 3E. In comparison to the pure 10% glycerol control, the laser beam in the lipidoid solution was more pronounced, indicating the presence of colloidal dissolved particles. Notably, the TE observed for the K16-PEG-N-PROXYL solution was enhanced in comparison to the other solutions of K16-PEG-S-PROXYL and the control PEG-N-PROXYL. This may result from either a higher particle concentration or a larger particle size.^23^

#### 2.2.2. Critical micelle concentration of model lipidoids

To further analyze the concentration dependent shift from a molecular to a colloidal solution, the CMC of the three compounds in 10% glycerol was analyzed using the pendant drop method to determine the surface tension as function of the lipidoid concentration. The results are shown in Figure 4. For PEG-N-PROXYL, the control compound without alkyl chains, no significant concentration dependent change in the surface tension was observed. The compound does not seem to be surface active and no CMC was determined in the screened concentration range. So most likely the aggregates and colloidal structures we observed through the DLS measurements and the Tyndall-effect experiments (2.2.1.) do not derive from classical micelles.

**Figure 4:**
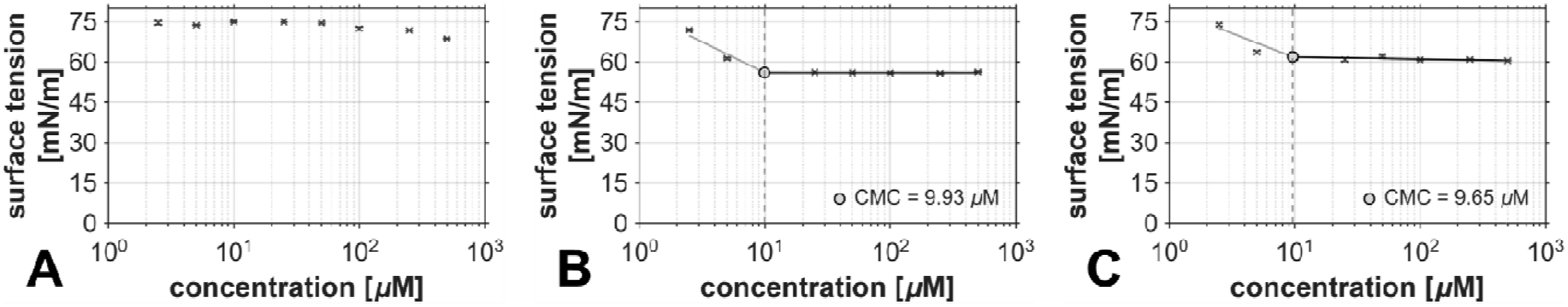
surface tension determined by the tensile drop method for different concentrations of (A) PEG-N-PROXYL; (B) K16-PEG-N-PROXYL; (C) K16-PEG-S-PROXYL. Measurements were performed in 10% glycerol. Results are given as mean and SD for three measurements (n = 3). CMC was determined by the intersection of two lines resulting in the biphasic graphs.

For K16-PEG-N-PROXYL and K16-PEG-S-PROXYL we observed a concentration dependent decrease of the surface tension until reaching a minimum. Both model lipidoids showed a comparable CMC of 9.9 µM and 9.7 µM. Consequently, at 50 µM (post-insertion conditions) both structures form micelles according to this experiment. Nevertheless, we did not detect particle sizes typical for micelles in the DLS at 50 µM (see chapter above). A small number of small micelles at this concentration (5times higher than CMC) let us assume that micelles were below detection limit of the DLS experiments.

Despite this self-aggregation, there still should be molecular dissolved lipids present for the post-insertion, since micelles are dynamic systems that reform constantly and are in equilibrium with dissolved molecules.^24^

Furthermore, the images of the droplets obtained during the CMC measurement (Figure S1) showed small structures for K16-PEG-N-PROXYL and K16-PEG-S-PROXYL at the highest concentration of 500 µM. These are very likely crystallites, which we possibly also detected in the DLS measurements at high concentration of the model lipidoids. However, the formation of these structures does not seem to affect the surface tension. The formation of these bigger aggregates could affect the post-insertion process in general, but in our post-insertion experiments the concentration of the model lipidoids was 50 µM, a concertation below the observed aggregate formation. So we concluded that the tendencies of the model lipidoids to form bigger agglomerates at higher concentrations should have no effect on our experiments.

### 2.3. Vesicle characterization

Since lipid composition of the target nanoparticle is another important parameter for the post-insertion of amphiphilic compounds in lipid-derived nanoparticles, we decided to use two different liposome compositions for the post-insertion experiments.^25^ 1,2-Distearoyl-*sn*-glycero-3-phosphocholine (DSPC) and cholesterol (Chol) are known to be main components of the outer shell of most LNP formulations.^26^ Therefore, we used DSPC:Chol (2:1) as one lipid formulation of interest. The phase transition temperature for DSPC is 55°C (gel phase L_β’_ at room temperature)^27^, nevertheless, the liquid ordered phase can be assumed to coexist for the investigated DSPC/Chol mixture.^28^ As “fluid” lipid formulation (alkyl chains with higher fraction of *gauche* conformation) we decided to use POPC with a reported phase transition temperature of −2°C (liquid crystalline L_α_ phase).^29^ Nevertheless, to not only rely on literature data, the phase transition in 10% glycerol was experimentally investigated via differential scanning calorimetry.

#### 2.3.1. Phase transition investigation via differential scanning calorimetry experiments

To verify that postulated phase behavior of the model LNP formulation is still present upon the addition of 10% glycerol to the solvent, differential scanning calorimetry (DSC) experiments of both liposome formulations were performed. Results are shown in Figure 5. The multilamellar liposomes of the two lipid compositions in water or 10% glycerol were prepared by using the dry lipid film hydration method, excluding the sonication procedure which would cause a decrease of particle size and lamellarity.

**Figure 5:**
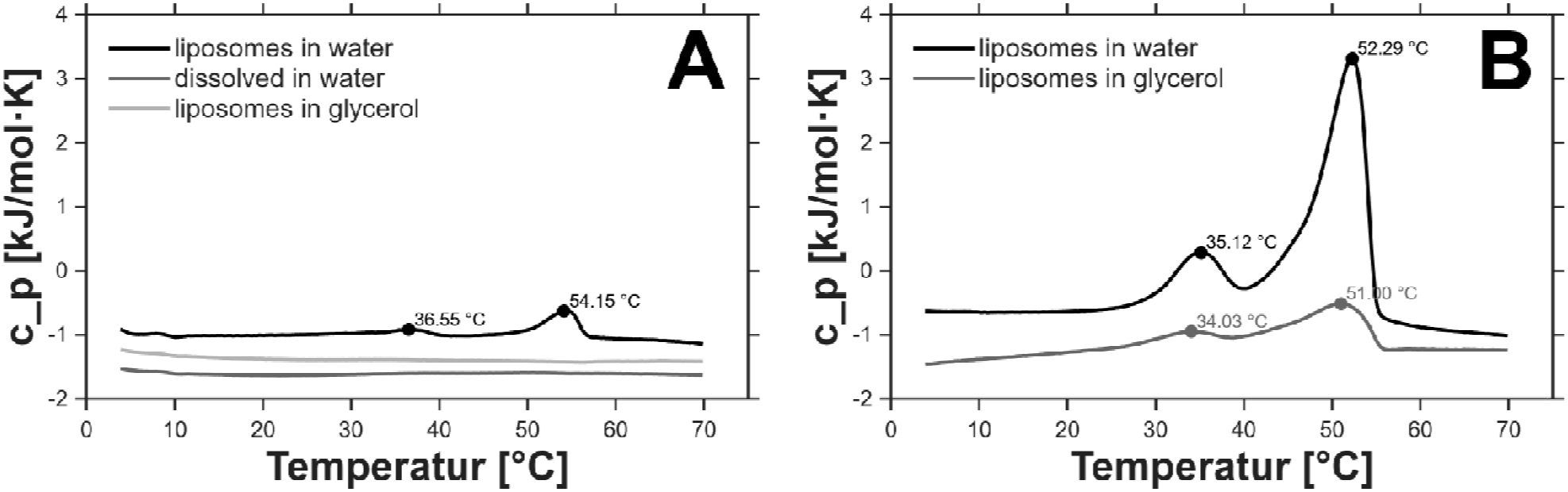
DSC heating scans of (A) POPC and (B) DSPC:Chol (2:1) liposomes in water and 10% glycerol. The phase transition temperature T_m_ is indicated as peak maximum of the scans. The scan labelled as dissolved in water in diagram A is the POPC sample where the powder was dispersed in water without using the thin film method for liposome preparation.

For the DSPC:Chol (2:1) liposomes, the phase transition temperatures in water and 10% glycerol were almost identical (see two identical peak maxima in Figure 5B). At room temperature the liposomes are in the gel phase or liquid ordered phase as assumed above. For the POPC liposomes, the DSC curve in 10% glycerol does not show a maximum of a phase transition in the scanned temperature range. This is in agreement with the reported phase transition temperature for POPC of −2°C, which is below the measured temperature range. We can assume the L_α_ phase.^29^ For POPC liposomes in water two small peaks with low intensity at 54 °C and 37°C were visible. Impurities can be excluded because the experiment was repeated several times while changing solvents. Hence, we assume this must be an effect of the formulation process. Nevertheless, POPC hydrolysis and the resulting gel phase stabilization caused by free fatty acids is an unlikely explanation, because we avoided sonication. Screening the literature identified effects of incomplete hydration of the POPC multilayers as possible explanation for the observed transition.^27^ To further investigate this, we decided to directly disperse POPC powder in water using short sonication, resulting in the absence of the weak transition at 54 °C.

#### 2.3.2. Liposome preparation for post-insertion experiments

Four different concentrations of the liposome formulations were prepared in 10% glycerol to analyze the post-insertion at four different model lipidoid-to-liposome ratios. To obtain liposomes with uniform size, sonication was followed by extrusion with a 100 nm membrane. The resulting liposomes were characterized by DLS to get information about the concentration dependence of the particle size (see Figure 6 and details in the supporting information Table S1). The DLS results indicated that this protocol yielded in POPC liposomes of consistent size in the range of 100 to 200 nm, even at the highest concentration (20 mg/mL). In contrast, the DPSC:Chol liposomes agglomerated after extrusion, resulting in increasing particle sizes up to 1100 nm at 20 mg/mL. Representative correlation functions and a *volume-*weighted size distribution curves for each liposome formulation are shown in Figures S2/S3. This difference in size could also influence the post-insertion process, because smaller liposomes can lead to a higher density of the post-inserted lipid.^30^ This is probably due to the higher membrane curvature. It is hard to estimate the effect of the increasing size observed for DSPC:Chol because it depends on the process which leads to the increased size. A fusion free agglomeration will not affect the insertion efficiency because the single liposomes will have the same curvature as the liposomes at the low concentration. An agglomeration with fusion results in bigger liposomes with less membrane curvature less effective for insertion of lipidoids.

**Figure 6:**
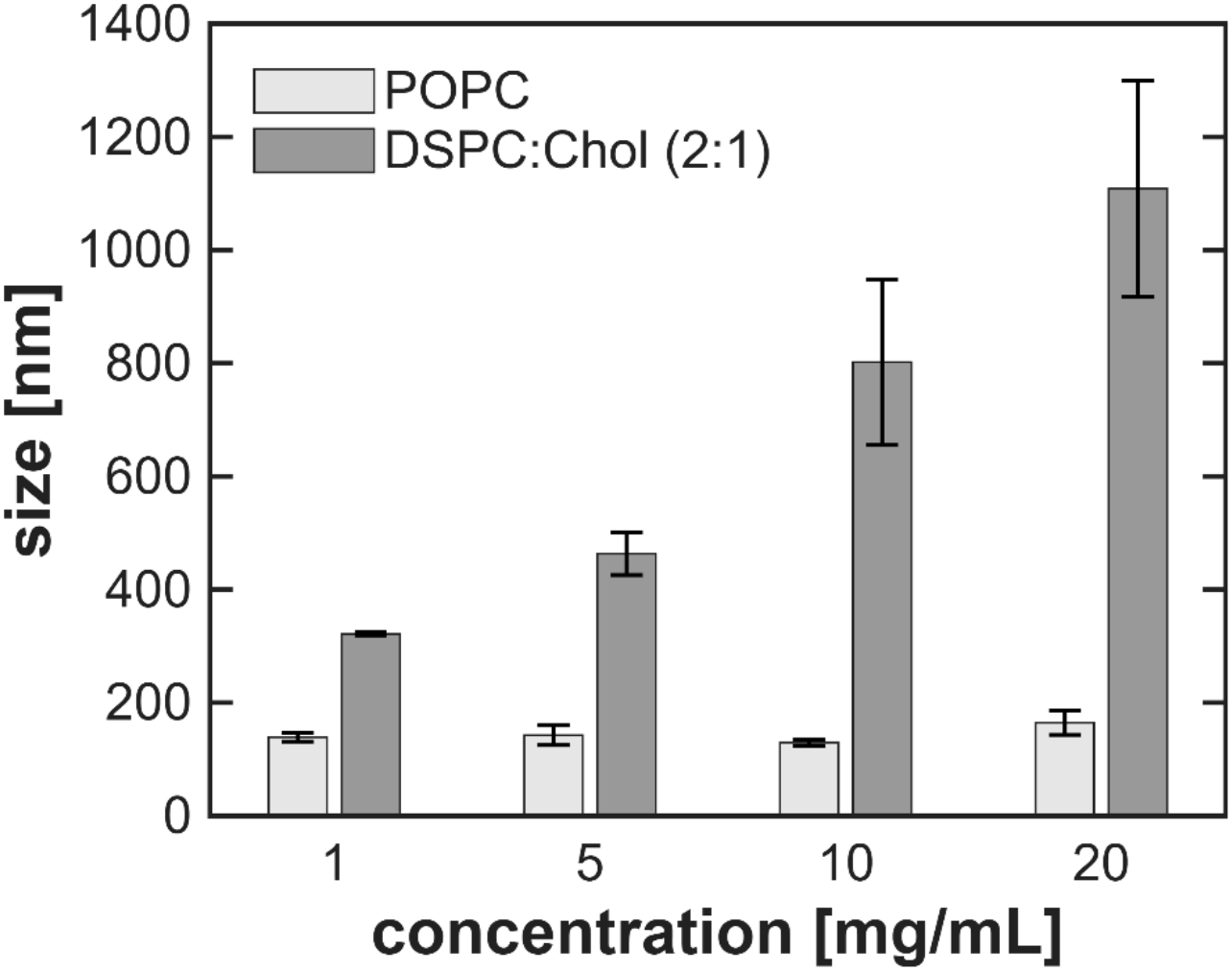
Particle size determined by DLS (z-average) of POPC and DSPC:Chol liposomes at different concentrations. Results are given as mean and SD of three measurements (n = 3).

### 2.4. Post-insertion characterization

We have chosen electron paramagnetic resonance (EPR) techniques to evaluate the interaction of the model-lipidoids with liposomes as lipid-based nanoparticles. To evaluate the amount of post-inserted model-lipidoids at different model lipidoid-to-liposome ratios, the post-insertion for each model lipidoid was performed for each liposomal formulation at four different liposome concentrations (1, 5, 10, and 20 mg/mL respectively). In this experimental setup, the concentration of each EPR-active model lipidoid was kept at 50 µM to achieve comparable signal intensities during EPR measurements.

#### 2.4.1. Continuous wave EPR experiments

CW-EPR is sensitive to the rotational mobility of the nitroxide label because the EPR spectrum reflects incomplete motional averaging of the anisotropic magnetic interactions between the unpaired electron and the external static magnetic field, namely the g-tensor and the nitrogen hyperfine coupling tensor.^31^ When the nitroxide undergoes rapid, isotropic reorientation on the microsecond timescale, as is the case for the free model lipidoids in 10% glycerol solution (Figure 7), these anisotropies are averaged and a sharp, three-line isotropic spectrum results. In contrast, insertion into a lipid bilayer restricts the label’s motion to axially symmetric tumbling about the long axis of the acyl chain, which incompletely averages the g and hyperfine interactions and produces the characteristic broad triplet “membrane” lineshape.^32^ This stark difference in spectral signature allows the two populations to be resolved within a single measurement, and their relative amounts can be determined from the linear combinations of the spectra of free and fully incorporated lipidoid. The results of these measurements are shown in Figure 7.

**Figure 7:**
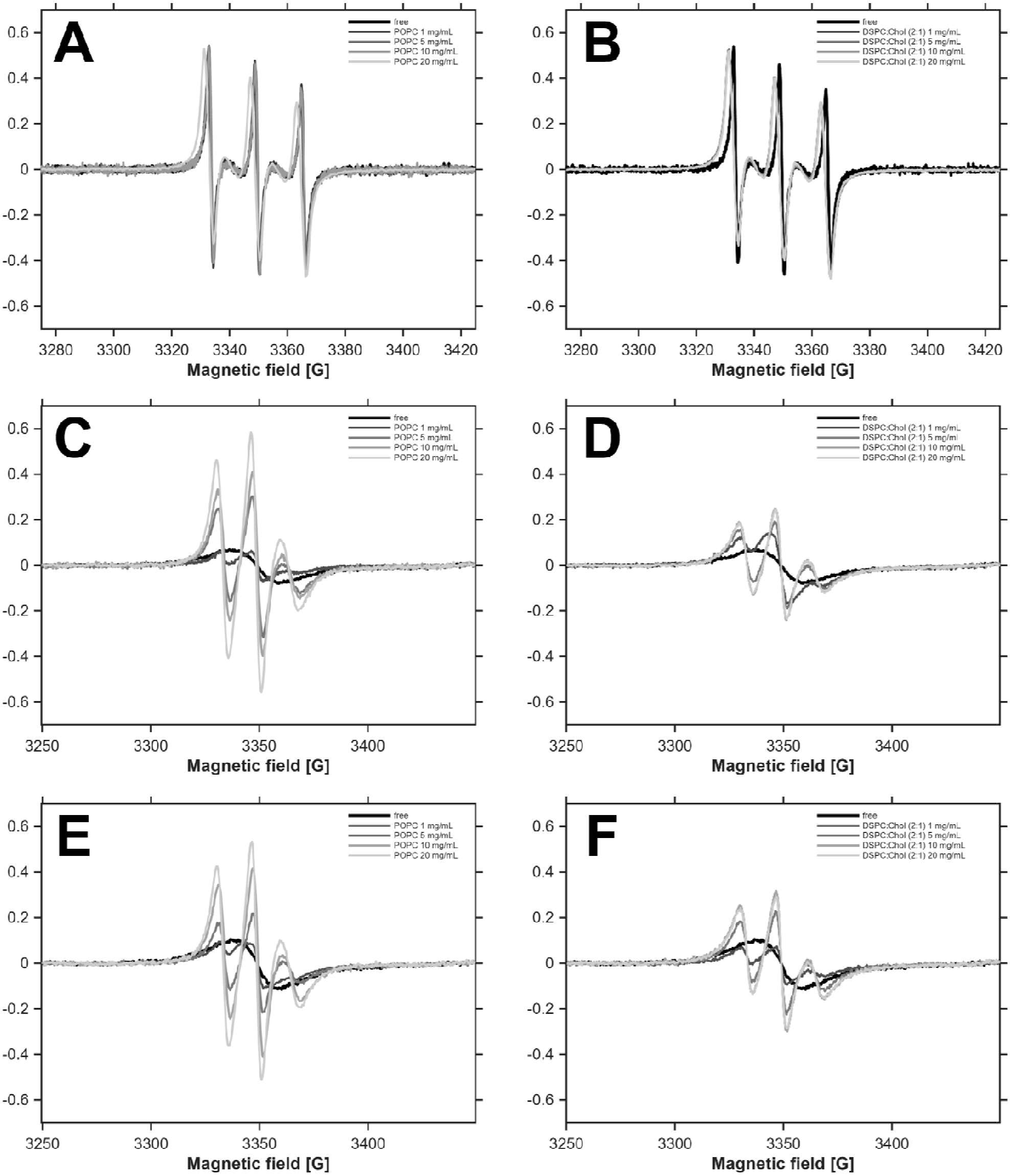
CW-EPR spectra of model lipidoids in 10 % glycerol (free) and after post-insertion in different concentrations of liposomes. (A) PEG-N-PROXYL and POPC liposomes; (B) PEG-N-PROXYL and DSPC:Chol (2:1) liposomes; (C) K16-PEG-N-PROXYL and POPC liposomes; (D) K16-PEG-N-PROXYL and DSPC:Chol (2:1); (E) K16-PEG-S-PROXYL and POPC; (F) K16-PEG-S-PROXYL and DSPC:Chol (2:1). Concentration of the model lipidoid for each measurement was 50 µM.

For PEG-N-PROXYL, the CW-EPR spectrum in absence of liposomes was identical to the spectra in presence of liposomes. Consequently, this reference model compound without alkyl chains did not interact with the lipid bilayer surfaces of liposomes for both liposomes with fluid or rigid alkyl chain packing. The compound showed no post-insertion tendencies and demonstrates that the lipophilic PROXYL groups do not introduce any unwanted interactions with liposome surface via integration. These results confirm the surface tension measurements which indicated no amphiphilic behavior (2.2.2.) and further supports the validity of PEG-N-PROXYL as negative control for post insertion.

For K16-PEG-N-PROXYL and K16-PEG-S-PROXYL, the spectrum of the model lipidoids in absence of liposomes was only one single, very broad line, see Figure 7 C-F. This indicates a strong exchange interaction of two close by spin systems occurring at distances below ∼10 Å, and suggests a self-aggregation of both acyl chains within individual free molecules.^33^ Upon addition of the liposomes, the typical hyperfine splitting was recovered, that became more pronounced with increasing liposome concentration. This result indicates the loss of short-distance exchange interactions within free model lipidoids upon incorporation into vesicles. Consequently, the binding of the model lipidoids to the liposome membrane resulted in a conformational change and a clearly distinguishable CW-EPR signal, compared to the signal of the model lipidoid free in solution. Additionally, the analysis of the isotropic hyperfine coupling constant indicated that the PROXYL groups are oriented to the aqueous medium after membrane insertion of the compound (see details chapter 5 Supporting Information).

To verify that the broad signal in absence of the liposomes is a result of a different conformation and not a result of micelle formation, CW-EPR spectra of the three compounds at a concentration of 5 µM were measured, which is below the CMC of K16-PEG-N-PROXYL and K16-PEG-S-PROXYL (2.2.2). Again, the spectra of the two model lipidoids were only single, broad lines, further indicating that the absence of hyperfine splitting is a result of self-aggregation of both acyl chains of individual free molecules. The respective spectra are shown in Figure S7.

To evaluate the efficiency of the post insertion of the model lipidoids into liposomal membranes, it was necessary to obtain reference spectra corresponding to the fully membrane-bound model lipidoids. For this purpose, the spectrum obtained after post-insertion at the highest liposome concentration (20 mg/mL) for each combination of model lipidoid and liposome formulation was corrected by subtracting the contribution of the spectrum of the model lipidoid freely in solution. This correction was performed iteratively until a symmetric spectrum with respect to the baseline was achieved, which was taken to represent the fully bound state. The determination of the amount of model lipidoid bound to liposome membrane in each sample, the contribution of the spectrum of the liposome free sample and the spectrum of the fully bound state (presence of 20 mg/mL liposomes) were determined by simulating a spectrum as a linear combination of the two that would overlap the measured spectrum. For illustration of this fitting process a representative example is presented in Figure 8 for K16-PEG-S-PROXYL after post-insertion in POPC liposomes. The fittings for the other samples can be found in the Supporting Information Figures S4-S6. The calculated contribution of the spectrum of the lipidoid free in solution (a) and the fully bound lipidoid (b) are shown in Table 1.

**Figure 8:**
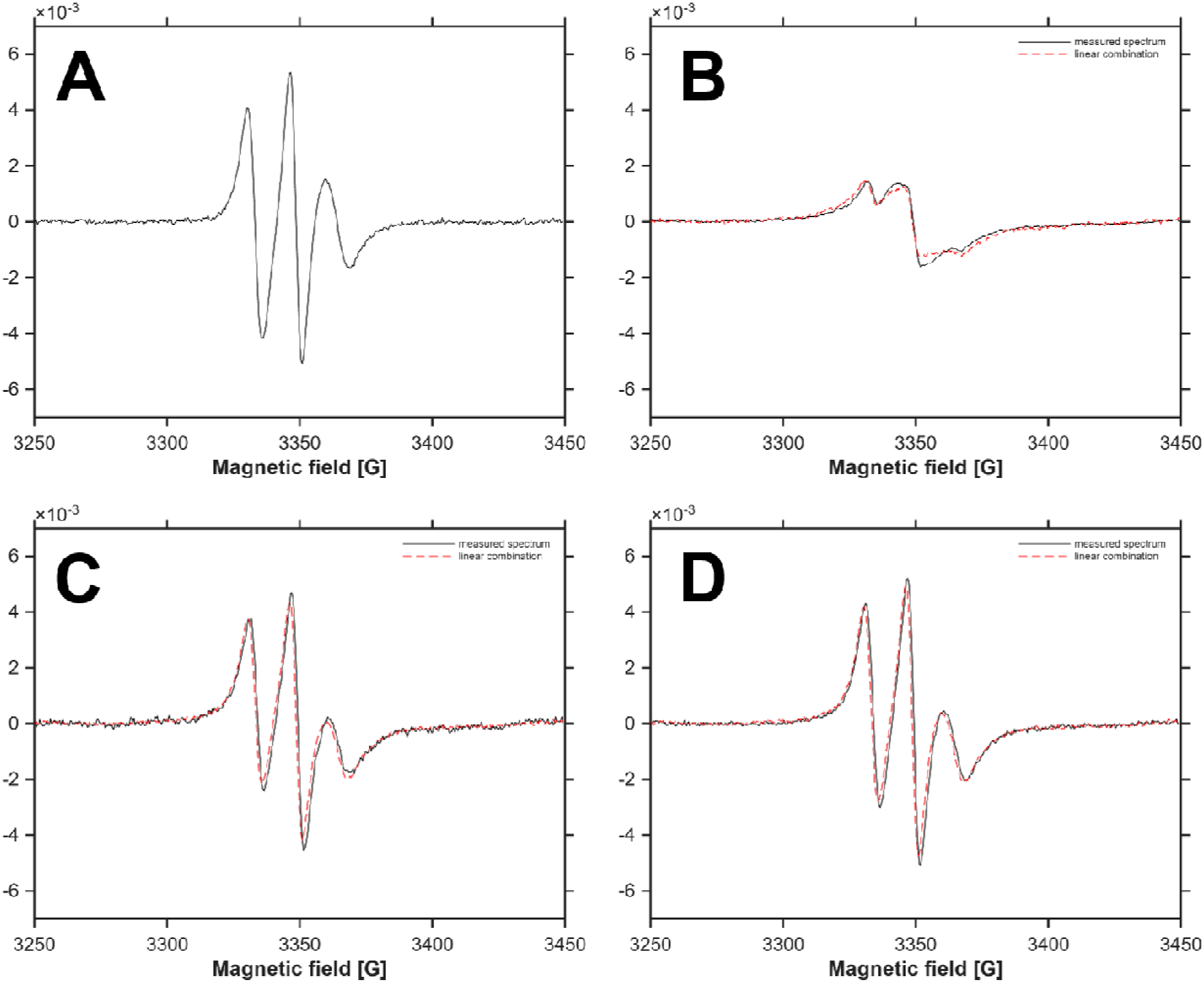
CW-EPR spectra of K16-PEG-S-PROXYL after post-insertion in POPC liposomes. (A) spectrum fully bound state. Measured spectrum (black) and calculated spectrum (red dotted line) as linear combination of the spectrum of the lipid free in solution and fully bound state after post insertion in (B) 1 mg/mL; (C) 5 mg/mL and (D) 10 mg/mL liposome.

**Table 1:**
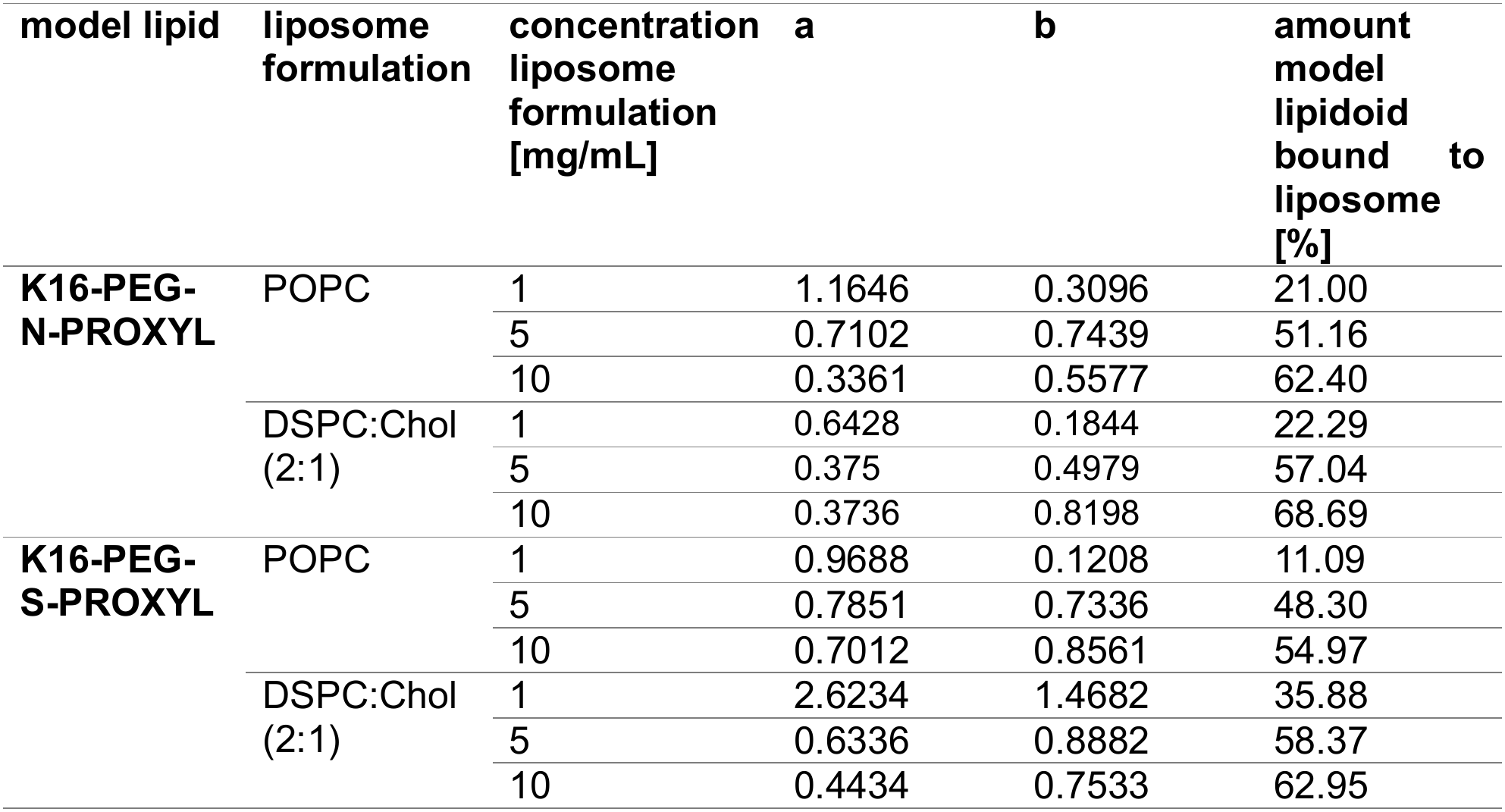
Amount of model lipidoid bound to liposome after post-insertion at different liposome concentrations. The amount was determined by calculation of the contribution of the CW spectrum of the model lipidoid free in solution (a) and the spectrum of the fully bound lipidoid (b) to the measured CW spectrum.

| **model lipid** | **liposome formulation** | **concentration liposome formulation [mg/mL]** | **a** | **b** | **amount model lipidoid bound to liposome [%]** |
|---|---|---|---|---|---|
| **K16-PEG-N-PROXYL** | POPC | 1 | 1.1646 | 0.3096 | 21.00 |
|  |  | 5 | 0.7102 | 0.7439 | 51.16 |
|  |  | 10 | 0.3361 | 0.5577 | 62.40 |
|  | DSPC:Chol (2:1) | 1 | 0.6428 | 0.1844 | 22.29 |
|  |  | 5 | 0.375 | 0.4979 | 57.04 |
|  |  | 10 | 0.3736 | 0.8198 | 68.69 |
| **K16-PEG-S-PROXYL** | POPC | 1 | 0.9688 | 0.1208 | 11.09 |
|  |  | 5 | 0.7851 | 0.7336 | 48.30 |
|  |  | 10 | 0.7012 | 0.8561 | 54.97 |
|  | DSPC:Chol (2:1) | 1 | 2.6234 | 1.4682 | 35.88 |
|  |  | 5 | 0.6336 | 0.8882 | 58.37 |
|  |  | 10 | 0.4434 | 0.7533 | 62.95 |

This data was used to model the lipidoid binding of the post-insertion process as Langmuir binding isotherm. Since the spectrum of the model lipidoid with 20 mg/mL liposomes was used to obtain the spectrum of the fully bound state, they were not included. The Langmuir model is used to describe the binding of substrates to surfaces and was therefore chosen as model of choice, knowing that it is a simplification of the interaction process connected with distinct requirements (homogeneous adsorption sites, individual binding of the molecule, dynamic reversible equilibrium, no interaction between molecules on the surface to alter their adsorption behavior)^34^. The percentage of model lipidoids in the initial post-insertion mixture was compared to the fraction of model lipidoids incorporated into the liposomes, relative to the total lipid content of liposomes. For post insertion it is desirable to employ a targeting lipid, that can be incorporated into the LNP at a high amount with high efficiency. Therefore, F_max_ and EC_50_ were determined using the Langmuir fit. The fits with the corresponding F_max_, EC_50_, and corresponding R^2^ values are summarized in Figure 9.

**Figure 9:**
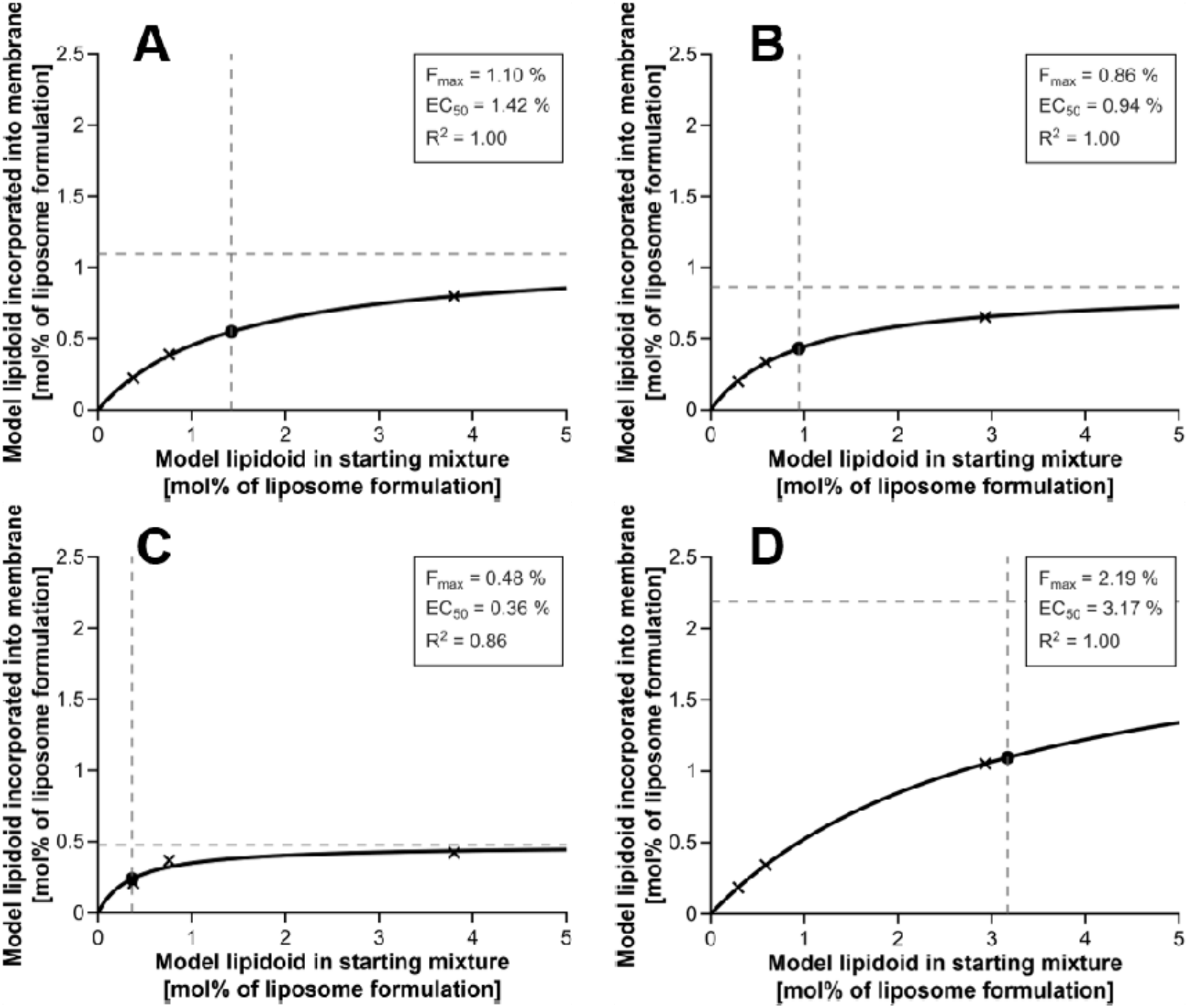
amount of model-lipidoids incorporated into liposome membrane at different model-lipidoid amounts in the post-insertion starting mixture. Fit according to the Langmuir model. F_max_, EC_50_ and R^2^ of the fit are shown. (A) K16-PEG-N-PROXYL and POPC liposomes; (B) K16-PEG-N-PROXYL and DSPC:Chol (2:1) liposomes; (C) K16-PEG-S-PROXYL and POPC liposomes; (D) K16-PEG-S-PROXYL and DSPC:Chol (2:1) liposomes

The determined F_max_ equals the maximum amount of model lipidoid, that can be post-inserted into the liposome formulation. This is relevant, because a higher amount of targeting-lipidoid is associated with higher cell uptake.^35^ For the POPC liposomes, K16-PEG-N-PROXYL had a higher F_max_ than K16-PEG-S-PROXYL (1.42 % and 0.48 %). In contrast, for the DSPC:Chol liposomes the F_max_ for K16-PEG-S-PROXYL was higher than K16-PEG-N-PROXYL (2.19 % and 0.86 %). This shows that for different LNP formulations a different design of the targeting lipidoids is necessary to achieve an optimal amount of targeting lipidoid. The determined EC_50_ equals the amount of lipidoid that has to be present in the post-insertion mixture at the beginning to achieve half of F_max_ in the resulting liposomes. The higher F_max_, the higher EC_50_ gets, meaning, that if a higher loading of the liposomes is possible, a higher amount of model lipidoid is needed to achieve this higher loading.

#### 2.4.2. Double electron-electron resonance experiments

A common EPR technique to determine interspin distance is double electron-electron resonance (DEER) which is based on the dipolar interaction between two paramagnetic centers. In case of our model lipidoids, DEER can be used to determine the intramolecular distance between the two PROXYL-labeled head groups and consequently the functionality of the lateral spacer. The resulting distance distributions are shown in Figure 10. The DEER measurements were performed for the model lipidoids and the control compound in 10% glycerol in absence of liposomes. Only for the control PEG-N-PROXYL we could obtain a distance distribution with a maximum at 16.8 Å (black curve Figure 10). For the two model lipidoids, the distance between the two spin probes was too small to be determined using DEER, which is limited to distances between 15 and 80 Å.^36^ This observation is in alignment with the results of the CW EPR measurements, which showed a strong exchange interaction, hinting at a short distance between the two spin probes. As described above, we assume that the free model lipidoids in solution form a loop, maybe driven by the alkyl chain interaction resulting in small distances between the PROXYL functionalities. Therefore, we performed the DEER measurements after post-insertion of the model-lipidoids at a concentration of 10 mg/mL POPC liposomes as matrix for insertion. In this experiments we could obtain a distance distribution in the range of around 15 to 35 Å. The mode of the distance distribution was 18.4 Å for K16-PEG-N-PROXYL and 17.7 Å for K16-PEG-S-PROXYL. This confirms that the binding of the model-lipidoids to the liposomal membrane leads to a conformational change of the model lipidoids, resulting in a stretching of the molecule and a higher distance of the two head groups and opening of the loop structure.

**Figure 10:**
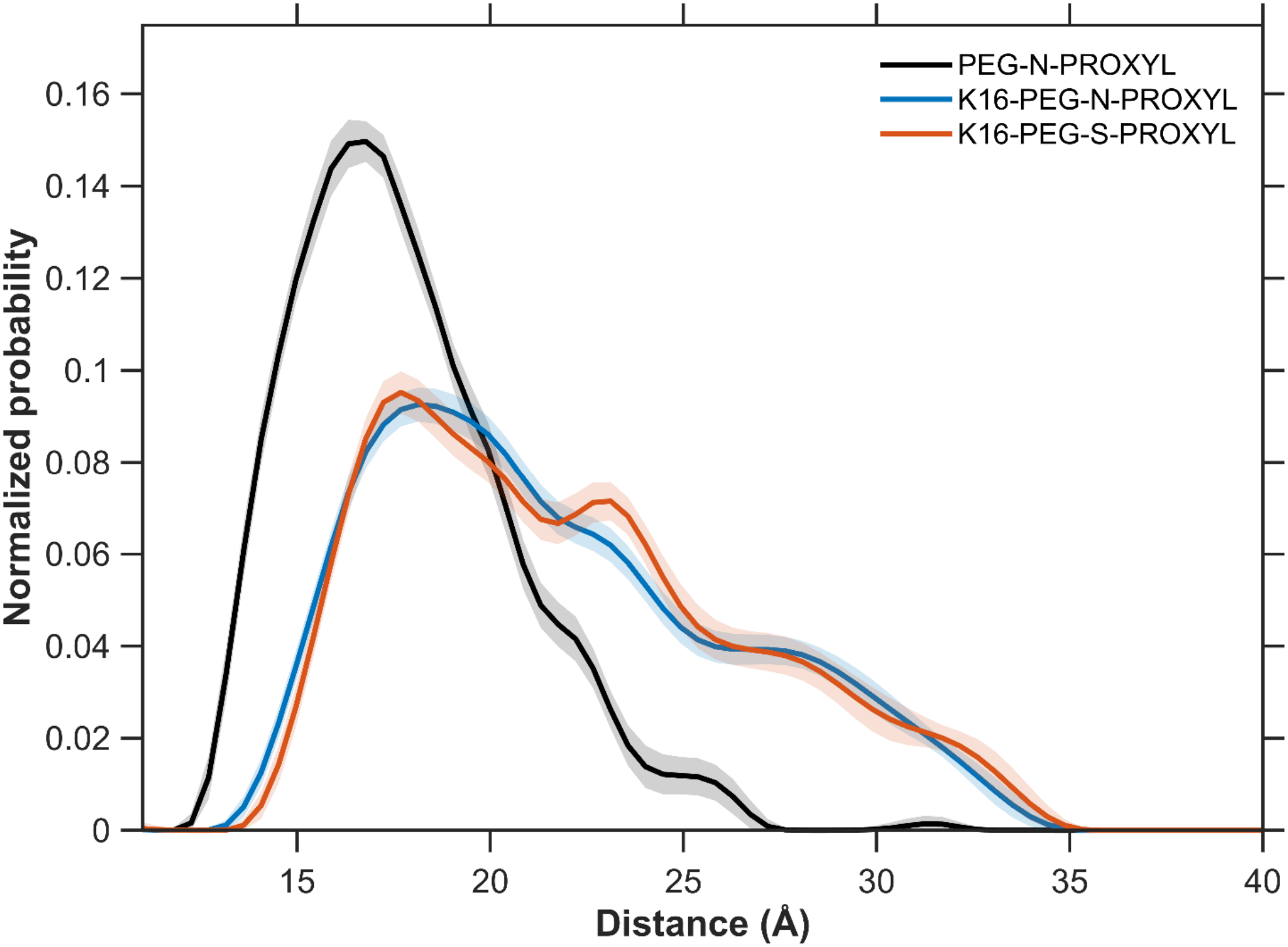
Overlays of normalized distance probability distributions of spin to spin distance of model lipids obtained by DEER analysis. DEER analysis was performed in 10 % glycerol for PEG-N-PROXYL and in 10% glycerol with 10 mg/mL POPC liposomes for K16-PEG-N-PROXYL and K16-PEG-S-PROXYL.

The experiment demonstrated that a certain lateral distance control between the two head groups of the bivalent model lipidoids is possible. However, this distance control was only present after incorporation of the lipidoids into the phospholipid layer of liposomes. The controlled distance was not locked ad a certain position, rather a broader distance distribution was observed. The mode of the distribution was for both lipidoids at a value around 18 Å (detailed values mentioned above), relatively comparable to the ≈17 Å measured for the soluble control compound. A pronounced difference was observed for the symmetry of the distance distribution. While the distribution curve is rather symmetric for the soluble control compound PEG-N-PROXYL, the distance distribution was positive skewed for both model lipidoids. The curves for K16-PEG-N-PROXYL and K16-PEG-S-PROXYL indicating a higher probability of the larger head group distances with valued above 20 Å. Consequently, the binding of the lipidoids at a lipid interface seem to allow other conformations than for a compound with lateral PEG spacer in aqueous solution. Oesterhelt et al. described that in aqueous environment PEG chains have a helical structure with the C-C-O-monomers in a *trans-trans-gauche* conformation resulting in a length of 2.8 Å per monomer.^37^ Energetically less favorable is the all-*trans* zigzag conformation leading to a fully stretched PEG chain with a length of 3.6 Å per monomer.^37^ This less favorable conformation seems to be (partially) present for the PEG-chains of the lipidoids at the liposome water interface resulting in the broad distribution with higher propability for the longer distances compared to the soluble derivative.

The wide range of the modelled distance between the two spins indicates a large conformational space, allowing a flexible distance between the two head groups of the lipid. This flexibility despite the controlled maximal distance could be a benefit for the binding of targeting moieties to their respective targets.

## 3. Conclusions

In this work we presented the synthesis and physico-chemical characterization of two model lipidoids with a lateral spacer for the separation of the head groups as a proof-of-concept for a dual targeting platform for the active targeting applicable on LNPs and liposomes. Using CW-EPR we were able to show the successful post-insertion of the two compounds into two different liposome formulations. The post-insertion efficiency was affected by different parameters: lipid species of the liposomes, chemical structure of the lipidoid to be inserted, and the model lipidoid-to-liposome ratio. Langmuir fits showed that the maximum amount of post-inserted lipidoid into the liposome formulation differs for each liposome/model lipidoid pairing. Interestingly, for the POPC liposomes K16-PEG-N-PROXYL had the higher F_max_, whereas K16-PEG-S-PROXYL had the higher F_max_ for the DSPC:Chol (2:1) liposomes. This shows that different liposome formulations need a different optimized targeting lipidoid for post-insertion.

The CW-EPR experiments also showed that the model lipidoids undergo a conformational change after binding to the liposome surface leading to a stretching of the molecule resulting in a longer distance. With DEER measurements we were able to verify this conformational change and proved the existence of a controlled distance between the two head groups. The large width of the DEER distance distributions indicate that the spin labels explore a large space only after incorporation in to nanoparticles. The benefit of this is twofold: (i) Free lipidoids will self-interact as loop structure preventing interference with functionalized LNP. (ii) Release of this self-interaction present for bound lipidoids allows large-scale dynamics likely required for efficiently exploring binding poses in target protein. Experiments investigating different attachment methods for the functional unit to access more rigid scaffolds are currently underway.

## 4. Materials and Methods

### 4.1. Materials

If not stated otherwise, all materials were obtained from Merck KGaA (Darmstadt, Germany). The PEG-Spacers 3,6,9,12,15-pentaoxaheptadecane-1,17-diamine and 4,7,10,13,16-pentaoxanonadecanedioic acid were obtained from BLD Pharmatech (Shanghai, China). The phospholipids 1-palmitoyl-2-oleoyl-*sn*-glycero-3-phosphocholine (POPC), 1,2-distearoyl-*sn*-glycero-3-phosphocholine (DSPC) as well as cholesterol were obtained from Avanti Polar Lipids (Alabaster, USA).

### 4.2. Synthesis

The synthesis of the model dual targeting lipidoids was adapted from established strategies in our group previously described for synthesis of lysine-based amino-functionalized lipids.^17^

The PROXYL labeled model lipidoids were characterized by ESI MS (LC-MS 2020 Shimadzu, Kyoto, Japan) (spectra shown in S8). ^1^H NMR was not possible for this compounds due to the PROXYL probe. Therefore, the precursor lipidoids of the EPR-active model lipidoids as well as the PEG-diamine were characterized by ^1^H NMR (Bruker AVANCE III HD 400, Bruker, Bruker corporation, Billerica, USA) (spectra shown in S9). Further retention factors (R_f_) for a given solvent after running ALUGRAM SIL G UV254 thin layer chromatography plates based on silica gel 60 (Macheray-Nagel). To evaluate substance spots UV light (154 nm) was used.

#### Synthesis of PROXYL-labeled PEG-spacer (compound 2 = PEG-N-PROXYL)

3,6,9,12,15-pentaoxaheptadecane-1,17-diamine (compound **1**) (0.25 mmol), 3-(carboxy)-2,2,5,5-tetramethyl-1-pyrrolidinyloxy (0.5 mmol), PyBob**^®^** (0.525 mmol) and DIPEA (1 mmol) were dissolved in CH_2_Cl_2_ (20 mL) and stirred under nitrogen atmosphere for 16 h. The solvent was evaporated on a rotary evaporator at reduced pressure. The crude compound was purified by column chromatography using silica gel 60 and CHCl_3_/MeOH (95:5, v/v).

3,6,9,12,15-pentaoxaheptadecane-1,17-diamine (compound **1**): C_12_H_28_N_2_O_5_, Yield – commercially acquired; R_f_ = 0.08 (CH_2_Cl_2_/MeOH, 9:1, v/v); ^1^H NMR (400 MHz, CD_3_OD) δ 3.66 (t, *J* = 4.1 Hz, 17H, O(C*H*_2_O)_4_); 3.55 (t, *J* = 5.3 Hz, 4H, OCH_2_C*H*_2_NH_2_); 2.82 (t, *J* = 5.3 Hz, 4H, C*H*_2_NH_2_)

Compound **2**, C_30_H_56_N_4_O_9_^2•^: Yield 81%; R_f_ = 0.18 (CHCl_3_/MeOH, 95/5, v/v) MS: *m/z:* 639.55 [M+Na] ^+^

#### Synthesis of double lysine-derivatized PEG-amin (compound 3)

3,6,9,12,15-pentaoxaheptadecane-1,17-diamine (compound **1**) (2 mmol), *N*-α-(9-fluorenylmethyloxycarbonyl)-*N*-ε-tert-butyloxycarbonyl-L-lysine (4 mmol), PyBob**^®^** (5 mmol) and DIPEA (8 mmol) were dissolved in CH_2_Cl_2_ (45 mL) and stirred under nitrogen atmosphere for 16 h. The solvent was evaporated on a rotary evaporator at reduced pressure. The crude compound was purified by column chromatography using silica gel 60 and CHCl_3_/MeOH (95:5, v/v).

#### Fmoc deprotection (compound 4)

The double lysine-derivatized PEG **3** (1.69 mmol) and piperidine (1.69 mmol) were dissolved in DMF (19 mL). The reaction was stirred for 3 h. The solvent was evaporated on a rotary evaporator at reduced pressure. The crude compound was purified by column chromatography using silica gel 60 and CHCl_3_/MeOH/25 % NH_3_ (9:1:0.2, v/v).

#### Coupling palmitic acid to double lysine-derivatized PEG (compound 5) and Boc deprotection (compound 6)

Compound **4** (0.95 mmol), palmitic acid (2.4 mmol), PyBob**^®^**(2.4 mmol), and DIPEA (3.8 mmol) were dissolved in CH_2_Cl_2_ and stirred under nitrogen atmosphere for 16 h. The solvent was evaporated in vacuum. The crude compound **5** was dissolved in HCl (37%, 1.8 mL) and CHCl_3_ (13 mL) and stirred for 2 h. Afterwards, ethanol (13 mL) was added to the solution. Ammonia solution (25%) was added dropwise under ice bath cooling until the pH shifted to 12. The organic layer was separated and extracted three times with CHCl_3_. The organic layers were combined, dried over Na_2_SO_4_, and the solvent was evaporated. The crude compound was purified by column chromatography using silica gel 60 and CHCl_3_/MeOH/25 % NH_3_ (8:2:0.2, v/v).

Compound **6**: C_56_H_112_N_6_O_9_; Yield: 41,5%; R_f_ = 0.52 (CHCl_3_/MeOH/NH_3_, 80/20/2, v/v/v); ^1^H NMR (400 MHz, CDCl_3_) δ 6.57 (d, *J* = 8.0 Hz, 2H), 4.51 (t, *J* = 7.0 Hz, 2H, COCH(N*H*CO)CH_2_), 3.69 – 3.66 (m, 9H) and 3.66 – 3.55 (m, 11H) and 3.55 – 3.47 (m, 2H) and 3.47 – 3.36 (m, 2H) NH(C*H*_2_C*H*_2_O)_5_C*H*_2_C*H*_2_NH; 2.74 (t, *J* = 6.7, 2.5 Hz, 4H, C*H*_2_NH_2_), 2.35 (s, 3H), 2.21 (t, *J* = 7.6 Hz, 4H, NOCC*H*_2_CH_2_), 1.90 – 1.77 (m, 2H) and 1.75 – 1.65 (m, 2H) and 1.62 (t, *J* = 6.6 Hz, 4H) and 1.57 – 1.46 (m, 4H) and 1.45 – 1.24 (m, 53H) NH_2_CH_2_(C*H*_2_)_3_, (C*H*_2_)_13_CH_3_, 0.89 (t, *J* = 6.7 Hz, 6H).

#### PROXYL labeling to yield model lipidoid K16-PEG-N-PROXYL

Compound **6** (0.1 mmol), 3-(carboxy)-2,2,5,5-tetramethyl-1-pyrrolidinyloxy (0.2 mmol), PyBob**^®^**(0.21 mmol) and DIPEA (0.4 mmol) were dissolved in CH_2_Cl_2_ (20 mL) and stirred under nitrogen atmosphere for 16 h. The solvent was evaporated in vacuum. The crude compound was purified by column chromatography using silica gel 60 and CHCl_3_/MeOH (95:5, v/v).

**K16-PEG-N-PROXYL**: C_74_H_140_N_8_O_13_^2•^: Yield 75%; R_f_ = 0.47 (CHCl_3_/MeOH, 95/5, v/v) MS: *m/z:* 1372.55 [M+Na] ^+^

The synthesis of the orthogonally protected lysine amide **9** and the deprotected lysine amide **10** have been described elsewhere.^17^

#### Coupling of lysine amide to PEG acid spacer (compound 12) and Boc deprotection to yield compound 13

The PEG-acid spacer **10** (1.5 mmol), the lysineamide **11** (3 mmol), PyBob**^®^** (3.15 mmol), and DIPEA (6 mmol) were dissolved in CH_2_Cl_2_ and stirred under nitrogen atmosphere for 16 h. The solvent was evaporated in vacuum. The crude compound **12** was dissolved in HCl (37%, 1.8 mL) and CHCl_3_ (13 mL) and stirred for 2 h. Afterwards, ethanol (13 mL) was added to the solution. Ammonia solution (25%) was added dropwise under ice bath cooling until the pH shifted to 12. The organic layer was separated and extracted three times with CHCl_3_. The organic layers were combined, dried over Na_2_SO_4_ and the solvent was evaporated. The crude compound was purified by column chromatography using silica gel 60 and CHCl_3_/MeOH/25 % NH_3_ (8:2:0.2, v/v).

Compound **13**: C_58_H_116_N_6_O_9_; Yield 42%, R_f_ = 41% (CHCl_3_/MeOH/NH_3_, 80/20/2, v/v/v); ^1^H NMR (400 MHz, CDCl_3_) δ 7.43 (d, *J* = 8.1 Hz, 2H), 6.92 (t, *J* = 5.7 Hz, 2H), 4.41 (t, *J* = 8.0, 5.6 Hz, 2H, COCH(N*H*CO)CH_2_), 3.81 – 3.72 (m, 4H) and 3.68 – 3.63 (m, 15H) NOC*H*_3_, NOCC*H*_2_(OC*H*_2_C*H*_2_)_4_C*H*_2_CON, 3.32 – 3.22 (m, 2H) and 3.16 (tdd, *J* = 13.1, 9.0, 5.3 Hz, 2H) NHC*H_2_*, 2.74 (t, *J* = 6.7 Hz, 4H, NH_2_C*H_2_*), 2.54 (t, *J* = 5.8 Hz, 5H), 2.48 (t, 4H, NOCC*H*_2_), 1.95 – 1.81 (m, 2H) and 1.68 (ddt, *J* = 13.6, 8.7, 4.1 Hz, 2H) and 1.60 –1.49 (m, 5H) and 1.49 – 1.37 (m, 7H) and 1.26 (d, *J* = 3.0 Hz, 52H) NH_2_CH_2_(C*H*_2_)_3_, (C*H*_2_)_13_CH_3_, 0.89 (t, *J* = 6.7 Hz, 6H, C*H*_3_).

#### PROXYL labeling to yield the model-lipidoid K16-PEG-S-PROXYL

Compound **13** (0.1 mmol), 3-(carboxy)-2,2,5,5-tetramethyl-1-pyrrolidinyloxy (0.2 mmol), PyBob**^®^**(0.21 mmol) and DIPEA (0.4 mmol) were dissolved in CH_2_Cl_2_ (20 mL) and stirred under nitrogen atmosphere for 16 h. The solvent was evaporated in vacuum. The crude compound was purified by column chromatography using silica gel 60 and CHCl_3_/MeOH (95:5, v/v).

***K16-PEG-S-PROXYL***: C_76_H_144_N_8_O_13_^2•^: Yield 43%; R_f_ = 0.04 (CHCl_3_/MeOH, 95/5, v/v) MS: *m/z:* 1401.00 [M+Na] ^+^

### 4.3. Determination Tyndall Effect

The sample was dissolved in 10% glycerol at a concentration of 1 mg/mL in a 1.5 mL flat bottom vial (VWR international, Radnor, USA). The solution was illuminated with a red laserpointer RF-LP-201 (conrad electronic SE, Hirschau, Germany). Pictures of the solution were taken with and without laser illumination.

### 4.4. Particle size measurements

Particle size measurements were carried out by dynamic light scattering (DLS) using a Litesizer 500 (Anton Paar GmbH, Graz-Straßgang, Austria) at 25°C with a measurement angle of 90° (side scatter) and a number of 10 runs. Measurements were performed in polystyrene cuvettes for the model lipidoid samples and quartz cuvettes for the liposome samples. The hydrodynamic diameter (z-average) and polydispersity index (PDI) were calculated by fitting the correlation function and applying the Stokes-Einstein equation using the Kalliope software. As viscosity 1.3 mPa*s was assumed for samples in 10% glycerol and 0.89 mPa*s for aqueous samples. The sample concentration for the liposome samples was adjusted at 20 µg/mL. Measurements of the model lipids were carried out at a sample concentration of 1 mg/mL and 50 µM respectively.

### 4.5. Critical micelle concentration (CMC) measurements

CMC determination was carried out by determining the surface tensions using the OCA 15 EC (Data Physics Instruments GmbH, Filderstadt, Germany). The surface tensions were calculated by using the pendant drop measurement method of the dpiMAX software. Drops were created using a DS 500/GT syringe and a SNS 050/026 needle with a dosing volume of 7 µL. The magnification was adjusted until the image of the drop was approximately 90% of the image and not changed during the measurements. The sample concentrations for the measurements were 2.5 µM, 5 µM, 10 µM, 25 µM, 50 µM, 100 µM, 250 µM, and 500 µM, respectively. The surface tension in dependence of concentration of the model lipidoids was graphically analyzed using MATLAB (R2024b, MathWorks, Natick, USA) and the CMC determined by the intersection of two tangents created in the graph.

### 4.6. Liposome preparation

Lipid stock solutions (10 mg/mL) of POPC and DSPC/cholesterol (2:1) were prepared in chloroform/methanol (8/2; v/v). The organic solvent was removed using a rotary evaporator at 500 mbar for 20 min, 400 mbar for 10 min, followed by drying the lipid film for 1 h at 5 mbar. 10% glycerol or water was added to the dried lipid films adhered to the glass wall to achieve a concentration of 1 mg/mL, 5 mg/mL, 10 mg/mL, or 20 mg/mL, respectively, and left overnight for hydratisation. The liposomes for the post-insertion experiments were afterwards sonicated for 30 min at 50 °C and 37 kHz using an Elmasonic P sonication bath (Elma Schmidbauer GmbH, Singen, Germany) and subsequently extruded 60 times using a 100 nm polycarbonate membrane. Liposome dispersions were stored at −8 °C.

### 4.7. Phase transition temperature (T_m_) measurements

The phase transition temperatures of the liposome dispersions (1 mg/mL) and POPC dispersion (2 mg/mL) were determined by differential scanning calorimetry using the PEAQ-DSC Cell Microcalorimeter (Malvern Panalytical, Kassel, Germany). The temperature range was scanned between 4°C and 70°C and the heating rate was 1 K/min. Each heating scan was repeated 3 times. Medium feedback was used for data recording. The reference cell was filled with either water or 10% glycerol, depending on the sample solvent. The water or 10% glycerol baseline was subtracted from the thermograms of the samples. The scans were analyzed using MATLAB (R2024b, MathWorks, Natick, USA), and the T_m_ determined by calculating the maximum of the heating scan.

### 4.8. Post-insertion

1.5 mM solutions of the model lipidoids in ethanol were prepared and added to 29 µL of the respective liposome dispersion with the resulting concentration of the model lipidoids being 50 µM. The Samples were incubated at room temperature for 1 h and afterwards stored at −8°C.

### 4.9. Continuous wave – electron paramagnetic resonance (CW-EPR) experiments

CW-EPR experiments were performed at room temperature on a EMXmicro with high sensitivity resonater (Bruker corporation, Billerica, USA). Spectral analysis was performed using MATLAB (R2024b, MathWorks, Natick, USA). All spectra were adjusted to the frequency used for the measurement of the respective model lipid in 10% glycerol and double integrated.

### 4.10. Double electron-electron resonance (DEER) experiments

DEER experiments were performed on a ElexSYS 580 (Bruker corporation, Billerica, USA). Spectral simulations and Data analysis were carried out using “Long Distances” (available under https://www.biochemistry.ucla.edu/Faculty/Hubbell/software.html).

### 4.11. Statistics and Langmuir fitting

The sample size of the statistically analyzed experiments is given as n below the corresponding figure. The results are given as mean ± standard deviation (SD). The statistically processing of the data was done using MATLAB (R2024b, MathWorks, Natick, USA). The binding data obtained by the CW-EPR measurements was analyzed using MATLAB (R2024b, MathWorks, Natick, USA). The data was fitted using the Langmuir fit to obtain F_max_ and EC_50_. The quality of the fit was assessed by calculating R^2^ (coefficient of determination). The results of the fits are given in the respective figure.

## Supporting information

Scheme 1 in high quality

## Acknowledgement

We acknowledge the Federal Ministry for Economic affairs and Energy (BMWE) for funding of the project ZellPro, Grant No. 16LP301102. The Graphical abstract and Figure 1 were created in BioRender. Kampik, V. (2026) https://BioRender.com/mmx4dka.

## Supplemental material

Additional data of the pendant drop method for CMC determination; additional data on the size evaluation of the model liposomes; additional data of the EPR measurements and evaluation; analytical details of the synthesized compounds; analysis of the isotropic hyperfine coupling constant

## Notes

### Competing Interest Statement

The authors have declared no competing interest.

## References

1. Barenholz, Y. (Chezy). Doxil® — The first FDA-approved nano-drug: Lessons learned. Journal of Controlled Release 160, 117–134 (2012).

2. Rachamala, H. K. Translational Advances in Lipid Nanoparticle Drug Delivery Systems for Cancer Therapy: Current Status and Future Horizons. Pharmaceutics 17, 1315 (2025).

3. Kang, D. D. et al. Targeting and tracking mRNA lipid nanoparticles at the particle, transcript and protein level. *Nat*. Biomed. Eng 9, 1591–1609 (2025).

4. Lin, Y., Cheng, Q. & Wei, T. Surface engineering of lipid nanoparticles: targeted nucleic acid delivery and beyond. swwlxb 9, 255 (2023).

5. Yoo, J., Park, C., Yi, G., Lee, D. & Koo, H. Active Targeting Strategies Using Biological Ligands for Nanoparticle Drug Delivery Systems. Cancers 11, 640 (2019).

6. Kroll, C. et al. Hybrid Bombesin Analogues: Combining an Agonist and an Antagonist in Defined Distances for Optimized Tumor Targeting. J. Am. Chem. Soc. 135, 16793–16796 (2013).

7. Veazey, R. S. et al. Dynamics of CCR5 Expression by CD4^+^ T Cells in Lymphoid Tissues during Simian Immunodeficiency Virus Infection. J Virol 74, 11001–11007 (2000).

8. Liu, J. et al. Low-Dose Mildronate-Derived Lipidoids for Efficient mRNA Vaccine Delivery with Minimal Inflammation Side Effects. ACS Nano 18, 23289–23300 (2024).

9. Patel, J. et al. Preparation of Targeted Lipid Nanoparticles by Insertion of Either Tetrazine–trans-Cyclooctene or Thiol–Maleimide Micelles. Bioconjugate Chem. 37, 406– 417 (2026).

10. Swart, L. E. et al. A robust post-insertion method for the preparation of targeted siRNA LNPs. International Journal of Pharmaceutics 620, 121741 (2022).

11. Mehrabian, A. et al. The comparison of biodistribution of glutathione PEGylated nanoliposomal doxorubicin formulations prepared by pre-insertion and post-insertion methods for brain delivery in normal mice. IET Nanobiotechnology 17, 112–124 (2023).

12. Xu, Y. et al. Lactoferrin-modified PEGylated liposomes loaded with doxorubicin for targeting delivery to hepatocellular carcinoma. IJN 5123 (2015) doi:10.2147/IJN.S87011.

13. Yazdian-Robati, R. et al. CD44-specific short peptide A6 boosts cellular uptake and anticancer efficacy of PEGylated liposomal doxorubicin in vitro and in vivo. Cancer Nano 14, 84 (2023).

14. Iden, D. L. & Allen, T. M. In vitro and in vivo comparison of immunoliposomes made by conventional coupling techniques with those made by a new post-insertion approach. Biochimica et Biophysica Acta (BBA) - Biomembranes 1513, 207–216 (2001).

15. Chen, C., Zhou, Y., Chen, C., Zhu, S. & Yan, X. Quantification of Available Ligand Density on the Surface of Targeted Liposomal Nanomedicines at the Single-Particle Level. ACS Nano 16, 6886–6897 (2022).

16. El-Gamal, F. R. et al. Synthesis and Evaluation of High Functionality and Quality Cell-penetrating Peptide Conjugated Lipid for Octaarginine Modified PEGylated Liposomes In U251 and U87 Glioma Cells. Journal of Pharmaceutical Sciences 111, 1719–1727 (2022).

17. Wölk, C. et al. General Synthesis and Physicochemical Characterisation of a Series of Peptide-Mimic Lysine-Based Amino-Functionalised Lipids. Chemistry A European J 19, 12824–12838 (2013).

18. Béduneau, A. et al. Design of targeted lipid nanocapsules by conjugation of whole antibodies and antibody Fab’ fragments. Biomaterials 28, 4978–4990 (2007).

19. Serrano, A. et al. Foxp3 inhibitory peptide encapsulated in a novel CD25-targeted nanoliposome promotes efficient tumor regression in mice. Acta Pharmacol Sin 46, 171– 183 (2025).

20. Zhang, Y. et al. Construction of a Targeting Nanoparticle of 3′,3″-Bis-Peptide-siRNA Conjugate/Mixed Lipid with Postinserted DSPE-PEG2000-cRGD. Mol. Pharmaceutics 16, 4920–4928 (2019).

21. Sugimoto, Y. et al. Microfluidic Post-Insertion Method for the Efficient Preparation of PEGylated Liposomes Using High Functionality and Quality Lipids. IJN **Volume** 17, 6675–6686 (2022).

22. Nag, O. K., Yadav, V. R., Hedrick, A. & Awasthi, V. Post-modification of preformed liposomes with novel non-phospholipid poly(ethylene glycol)-conjugated hexadecylcarbamoylmethyl hexadecanoic acid for enhanced circulation persistence in vivo. International Journal of Pharmaceutics 446, 119–129 (2013).

23. Xiao, W. et al. Highly Sensitive Colorimetric Detection of a Variety of Analytes via the Tyndall Effect. Anal. Chem. 91, 15114–15122 (2019).

24. Guo, W. et al. Micelle Dynamic Reconstruction to Effectively Modulate the Transmission of Smart Windows. J. Phys. Chem. B 126, 4872–4880 (2022).

25. Golkar, N., Tamaddon, A. M. & Samani, S. M. Effect of lipid composition on incorporation of trastuzumab-PEG-lipid into nanoliposomes by post-insertion method: physicochemical and cellular characterization. Journal of Liposome Research 1–13 (2015) doi:10.3109/08982104.2015.1048692.

26. Trollmann, M. F. W. & Böckmann, R. A. mRNA lipid nanoparticle phase transition. Biophysical Journal 121, 3927–3939 (2022).

27. Chen, W., Duša, F., Witos, J., Ruokonen, S.-K. & Wiedmer, S. K. Determination of the Main Phase Transition Temperature of Phospholipids by Nanoplasmonic Sensing. Sci Rep 8, 14815 (2018).

28. Tamai, N. et al. A new interpretation of eutectic behavior for distearoylphosphatidylcholine–cholesterol binary bilayer membrane. Biophysical Chemistry 135, 95–101 (2008).

29. Żak, A., Korshunova, K., Rajtar, N., Kulig, W. & Kepczynski, M. Deciphering Lipid Arrangement in Phosphatidylserine/Phosphatidylcholine Mixed Membranes: Simulations and Experiments. Langmuir 39, 18995–19007 (2023).

30. Lee, S.-H., Sato, Y., Hyodo, M. & Harashima, H. Size-Dependency of the Surface Ligand Density of Liposomes Prepared by Post-insertion. Biological & Pharmaceutical Bulletin 40, 1002–1009 (2017).

31. Gaffney, B. J. [17] Spin-label measurements in membranes. in Methods in Enzymology vol. 32 161–198 (Elsevier, 1974).

32. McConnell, H. M. & Hubbell, W. L. Molecular motion in spin-labeled phospholipids and membranes. J. Am. Chem. Soc. 93, 314–326 (1971).

33. Eaton, S. S., Woodcock, L. B. & Eaton, G. R. Continuous wave electron paramagnetic resonance of nitroxide biradicals in fluid solution. Concepts Magnetic Resonance **47A**, e21426 (2018).

34. Latour, R. A. The langmuir isotherm: A commonly applied but misleading approach for the analysis of protein adsorption behavior. J Biomedical Materials Res 103, 949–958 (2015).

35. Re, F. et al. Functionalization of liposomes with ApoE-derived peptides at different density affects cellular uptake and drug transport across a blood-brain barrier model. Nanomedicine: Nanotechnology, Biology and Medicine 7, 551–559 (2011).

36. Peter, M. F. et al. Cross-validation of distance measurements in proteins by PELDOR/DEER and single-molecule FRET. Nat Commun 13, 4396 (2022).

37. Oesterhelt, F., Rief, M. & Gaub, H. E. Single molecule force spectroscopy by AFM indicates helical structure of poly(ethylene-glycol) in water. New J. Phys. 1, 6–6 (1999).

