## supplement material for "Design of colipids with controlled lateral distance of head group moieties for targeting strategies in the field of lipid-based nanoparticles"

**Content:**

1. **Additional data of the pendant drop method for CMC determination**
2. **Additional data on the size evaluation of the model liposomes**
3. **Additional data of the EPR measurements and evaluation**
4. **Analytical details of the synthesized compounds**
5. **Analysis of the isotropic hyperfine coupling constant**

### 1. Additional data of the pendant drop method for CMC determination

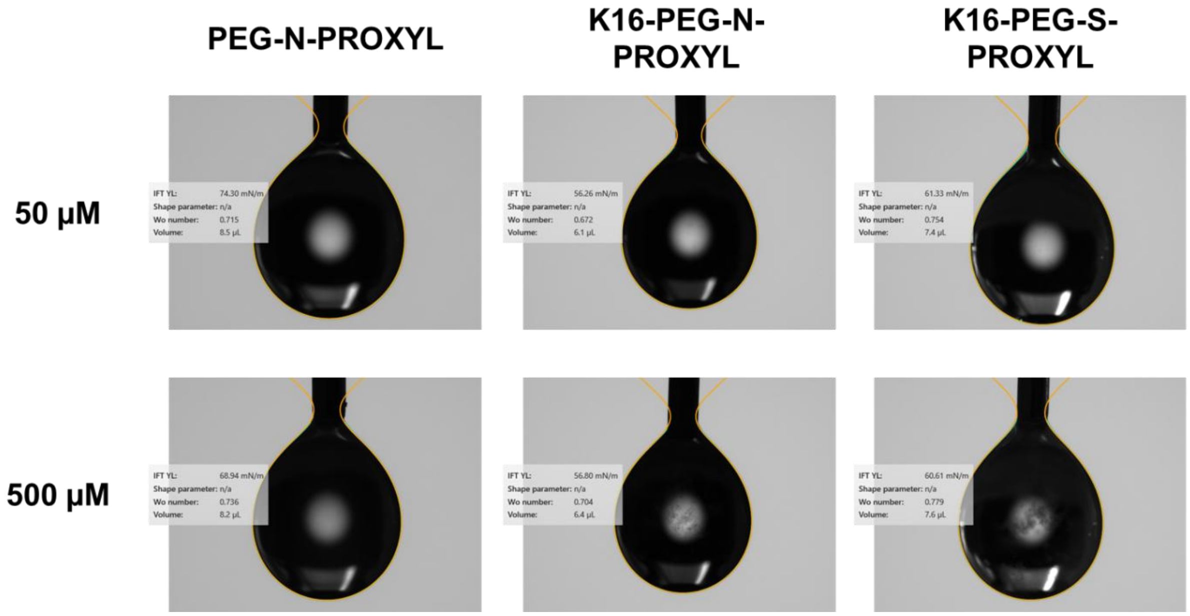


**Figure S1:** exemplary images of droplets of PEG-N-PROXYL, K16-PEG-N-PROXYL and K16-PEG-S-PROXYL at 50 µM and 500 µM in 10% Glycerol

### 2. Additional data on the size evaluation of the model liposomes

**Table S1:** particle size (z-average) and poly dispersity index (PDI) of POPC and DSPC:Chol (2:1) liposomes at different concentrations. Particle size and PDI were determined using DLS. Results are given as mean and SD of three measurements (n = 3).

| | **POPC** | | **DSPC/Chol (2:1)** | |
|---|---|---|---|---|
| **concentration [mg/ml]** | hydrodynamic diameter [nm] | PDI [%] | hydrodynamic diameter [nm] | PDI [%] |
| **1** | 138.78 +/- 7.86 | 24.3 +/- 1.7 | 321.3 +/- 2.85 | 27.6 +/- 2.3 |
| **5** | 142.45 +/- 17.7 | 17.7 +/- 1.8 | 463.4 +/- 37.8 | 29.7 +/- 1.7 |
| **10** | 129.34 +/-5.49 | 26.8 +/- 3.2 | 802 +/- 146.41 | 30.4 +/-2.8 |
| **20** | 164.58 +/- 21.64 | 26.9 +/- 2.7 | 1108.8 +/- 190.87 | 30.1 +/- 3.7 |

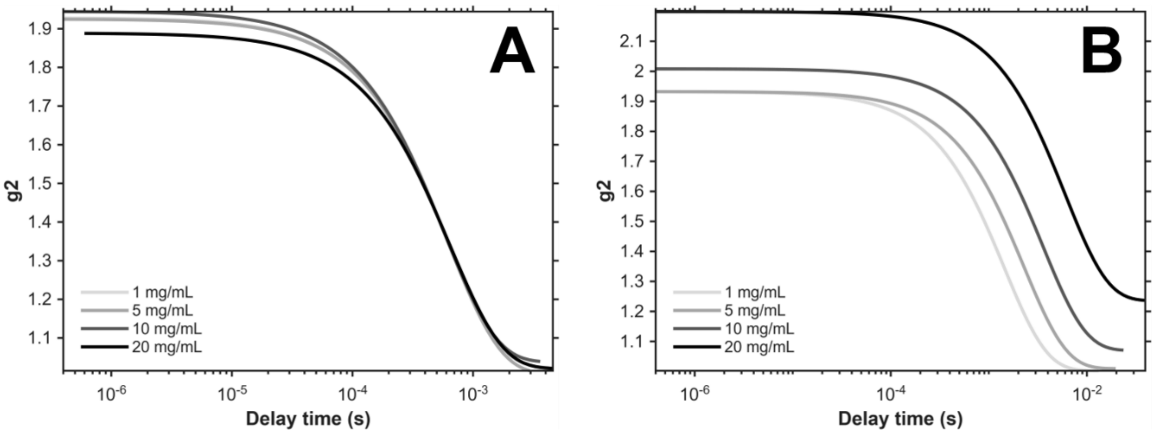


*Figure S2: correlation function of DLS measurements at 1, 5, 10 and 20 mg/mL of (A) POPC Liposomes and (B) DSPC:Chol (2:1) liposomes. All measurements have been performed in 10% glycerol.*

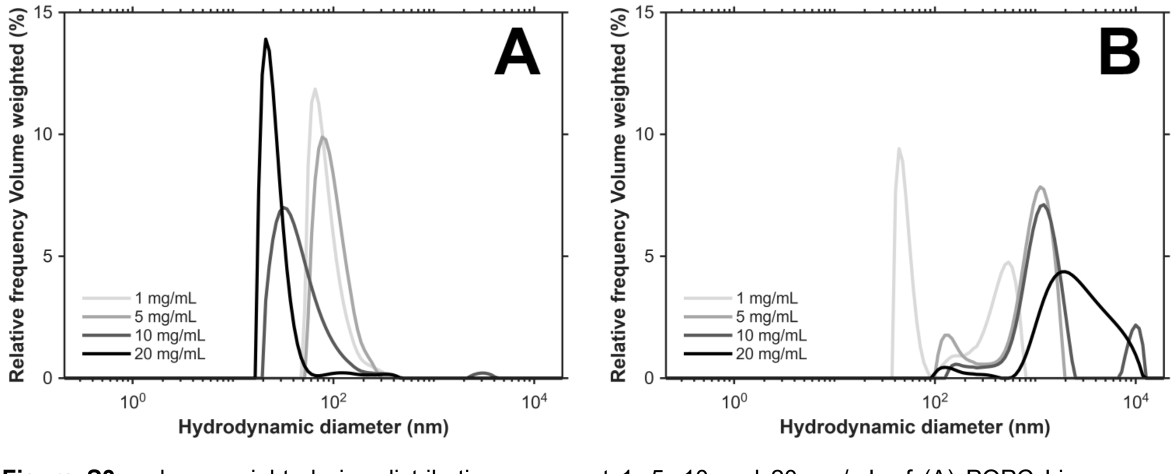


**Figure S3:** volume-weighted size distribution curves at 1, 5, 10 and 20 mg/mL of (A) POPC Liposomes and (B) DSPC:Chol (2:1) liposomes. All measurements have been performed in 10% glycerol.

### 3. Additional data of the EPR measurements and evaluation

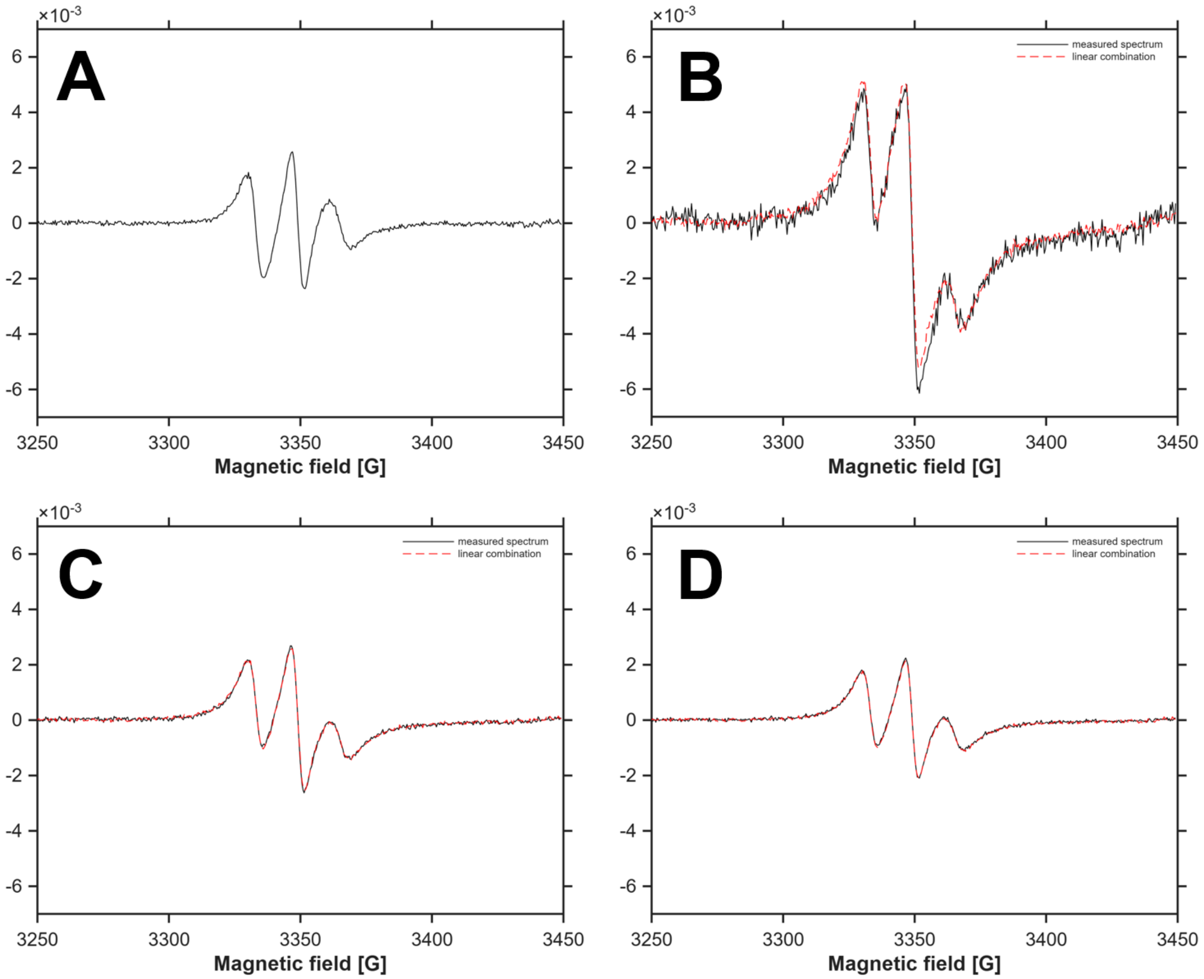


**Figure S4:** CW-EPR spectra of K16-PEG-S-PROXYL after post-insertion in DSPC:Chol (2:1) liposomes. (A) spectrum fully bound state. Measured spectrum (black) and calculated spectrum as linear combination of the spectrum of the lipid free in solution and fully bound state after post insertion in (B) 1 mg/ml; (C) 5 mg/ml and (D) 10 mg/ml liposome.

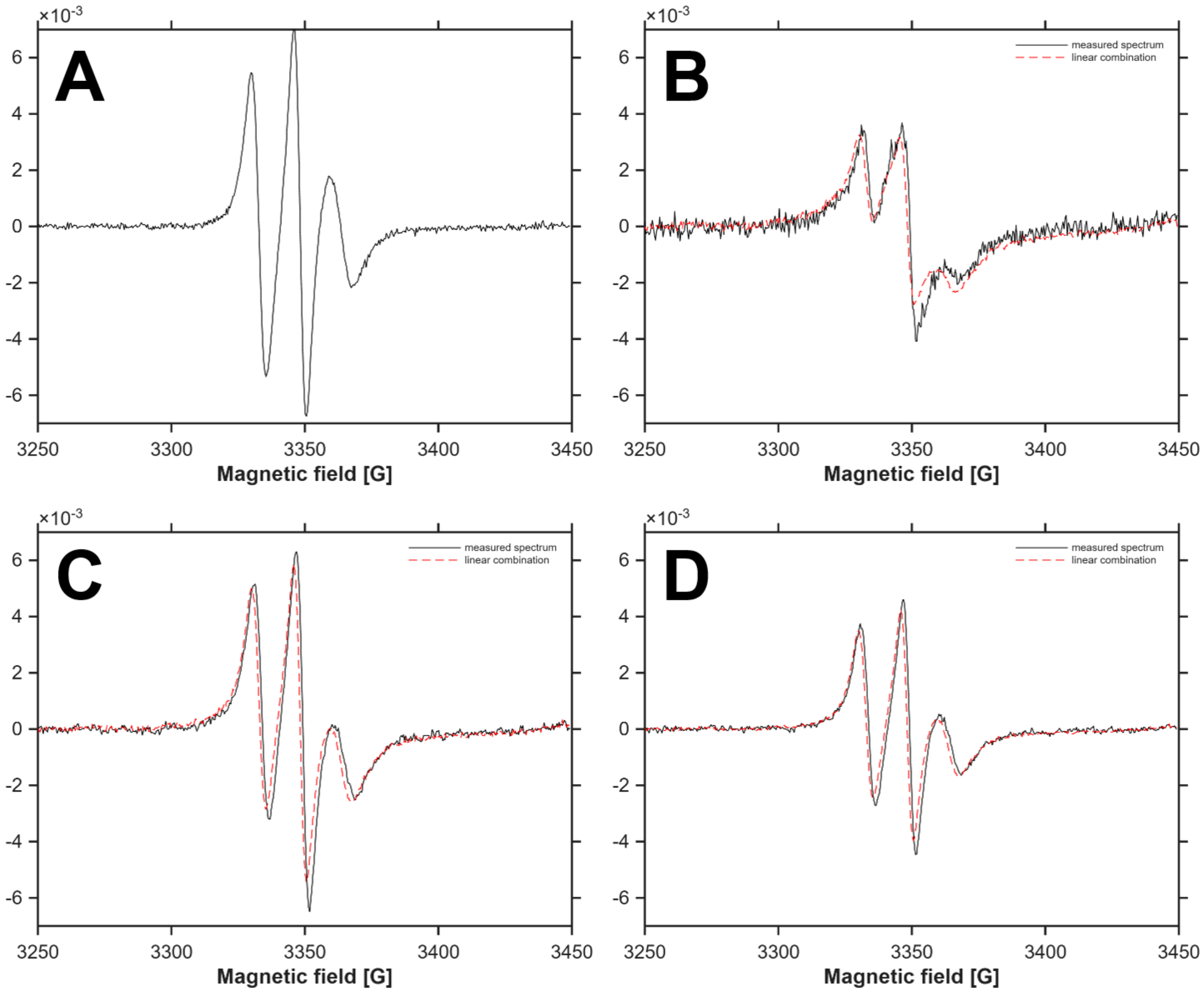


**Figure S5:** CW-EPR spectra of K16-PEG-N-PROXYL after post-insertion in POPC liposomes. (A) spectrum fully bound state. Measured spectrum (black) and calculated spectrum as linear combination of the spectrum of the lipid free in solution and fully bound state after post insertion in (B) 1 mg/ml; (C) 5 mg/ml and (D) 10 mg/ml liposome.

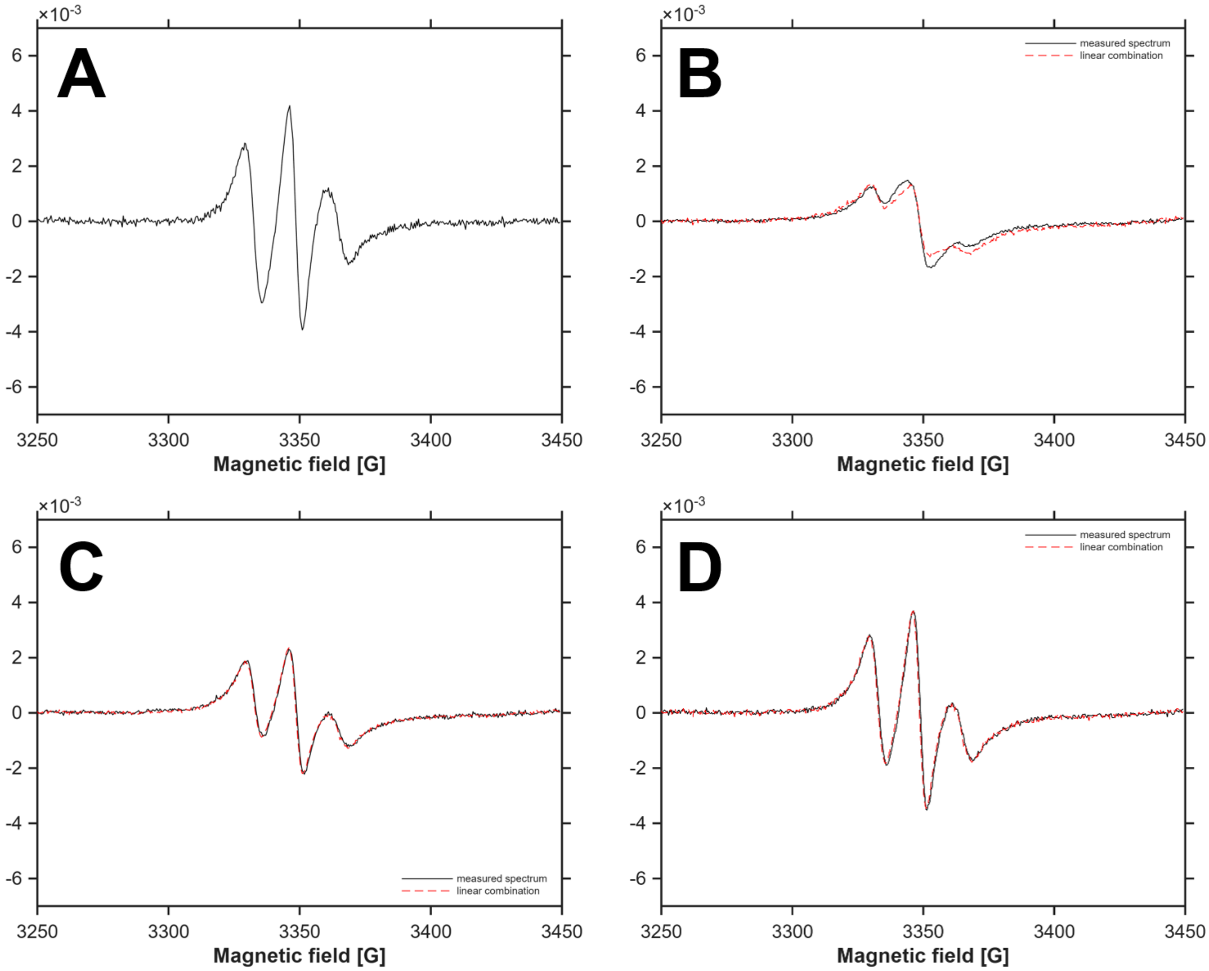


**Figure S6:** CW-EPR spectra of K16-PEG-N-PROXYL after post-insertion in DSPC:Chol (2:1) liposomes. (A) spectrum fully bound state. Measured spectrum (black) and calculated spectrum as linear combination of the spectrum of the lipid free in solution and fully bound state after post insertion in (B) 1 mg/ml; (C) 5 mg/ml and (D) 10 mg/ml liposome.

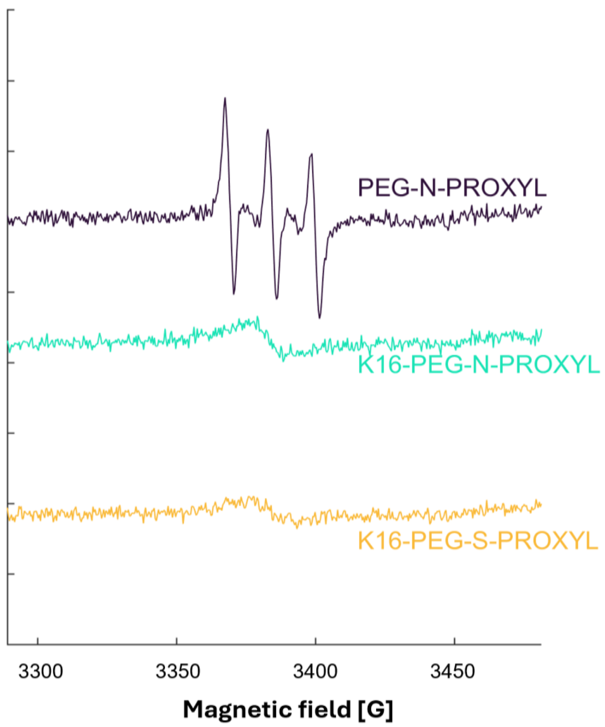


**Figure S7:** CW-EPR spectra PEG-N-PROXYL, K16-PEG-N-PROXYL and K16-PEG-S-PROXYL at 5 µM

### 4. Analytical details of the synthesized compounds

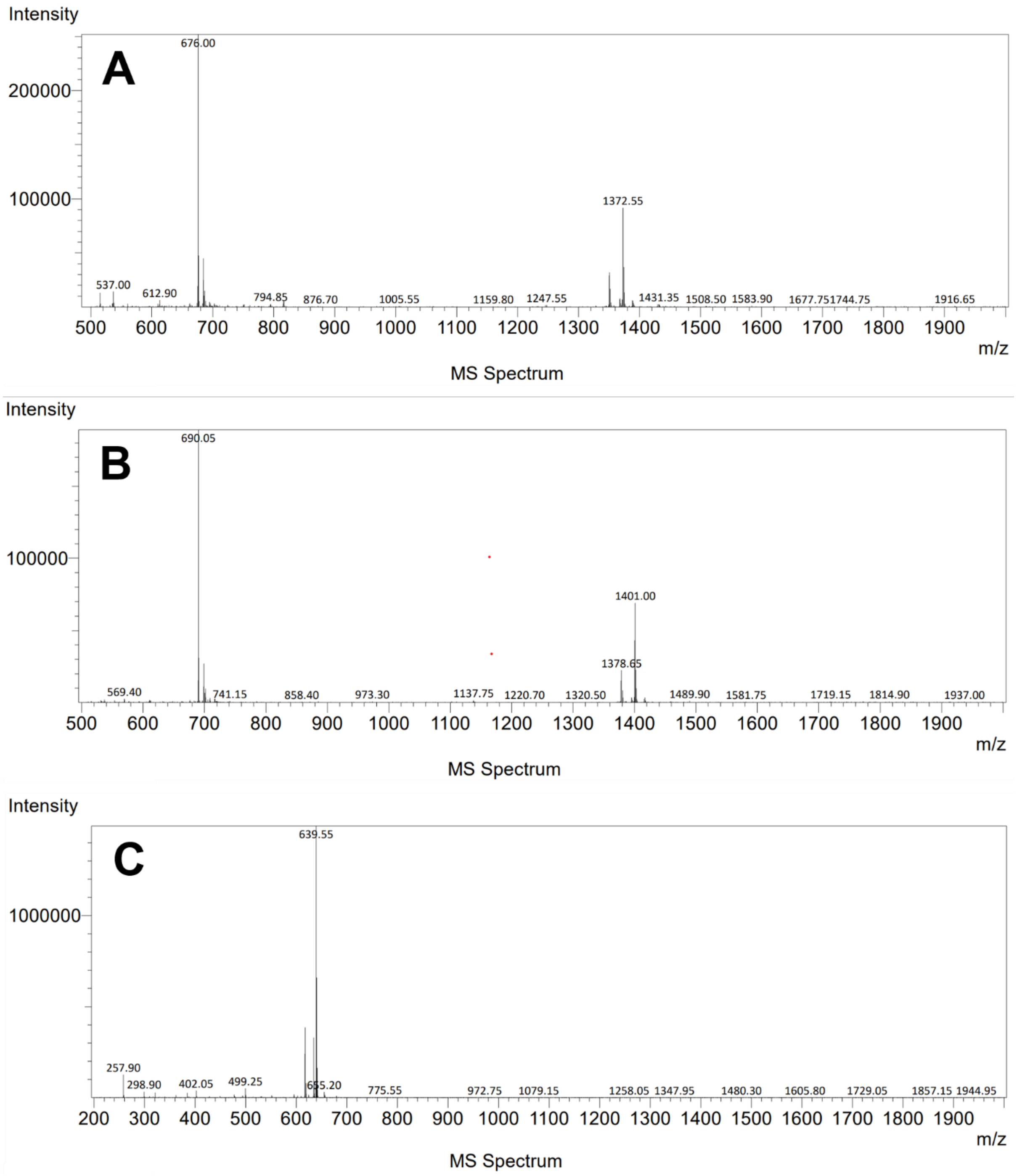


**Figure S8:** mass spectra of (A) K16-PEG-N-PROXYL, (B) K16-PEG-S-PROXYL, and (C) PEG-N-PROXYL

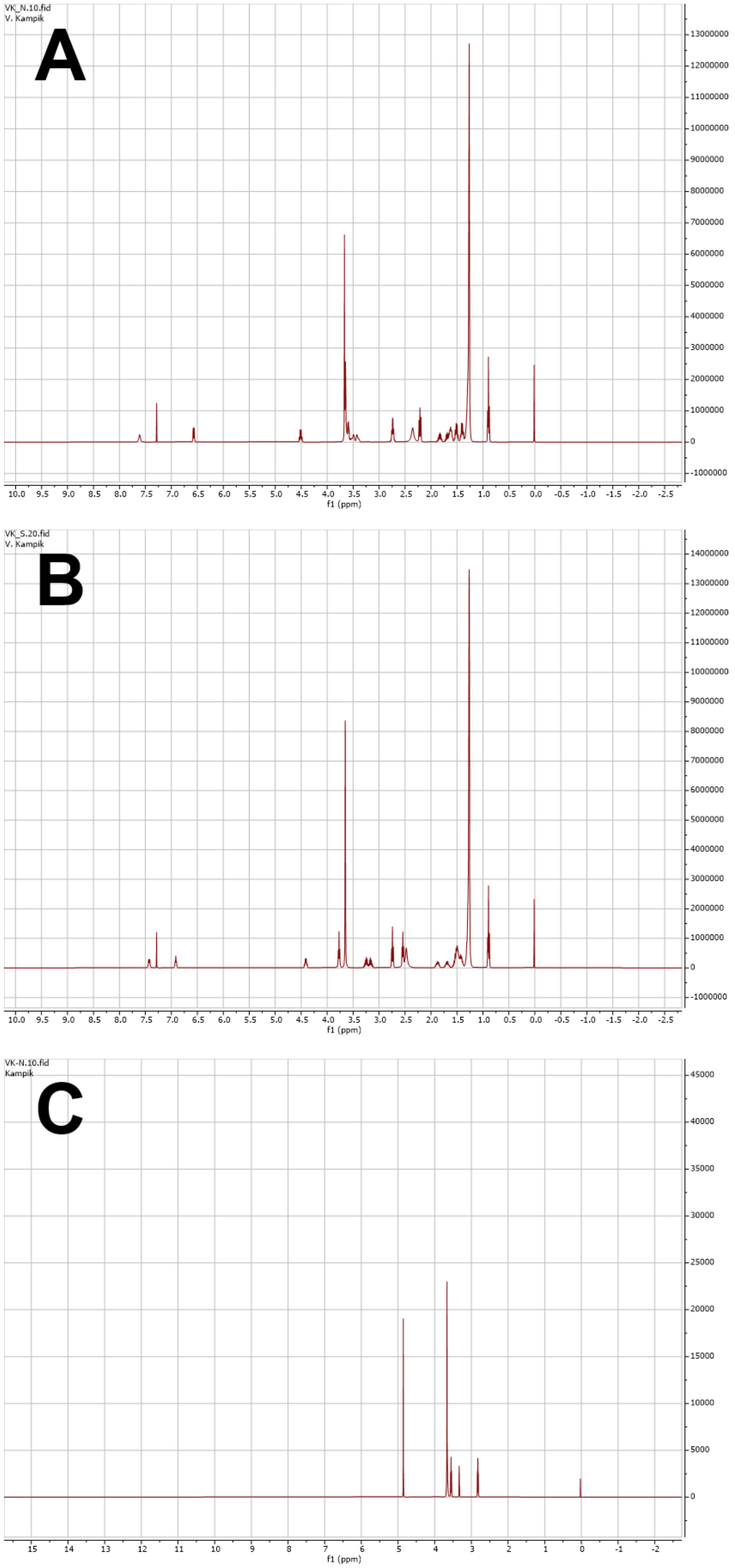


**Figure S9:** NMR spectra of (A) compound **6**, (B) compound **13**, and (C) compound **1**

### 5. Analysis of the isotropic hyperfine coupling constant

To test whether the PROXYL headgroup remains exposed to the aqueous phase after the model lipidoids bind to the liposomes, we determined the isotropic $^{14}N$ hyperfine coupling constant, $A_N$. $A_N$ decreases when a nitroxide partitions from water into the hydrocarbon region of a lipid bilayer (Hubbell & McConnell, 1968), and it tracks the transmembrane polarity profile, decreasing toward the bilayer center (Marsh, 2001). We obtained $A_N$ as half the field separation between the low-field and high-field lines of the three-line spectrum. Spectra were evaluated for the lipidoids fully bound to POPC and to DSPC/Chol liposomes (Figure 8 of the main text; Figures S4-S6). The free lipidoids in solution could not be evaluated because of strong spin-exchange broadening. As a reference for a water-exposed PROXYL label, we therefore used the unbound control compound PEG-N-PROXYL in 10% glycerol (Figure 7A/B; Figure S7). For both model lipidoids, $A_N$ in the liposome-bound state agrees with that of the unbound control compound within experimental uncertainty (Table S2). This indicates that the PROXYL headgroup does not penetrate the hydrocarbon region of the bilayer but remains in the polar interfacial region, exposed to the surrounding aqueous phase.

**Table S2**: $A_N$ of PROXYL label of PEG-N-PROXYL, K16-PEG-N-PROXYL and K16-PEG-S-PROXYL unbound in 10% Glycerol, fully bound to POPC liposomes and fully bound to DSPC/Chol liposomes

| | **Unbound in 10% glycerol** | **Fully bound to POPC liposomes** | **Fully bound to DSPC/Chol liposomes** |
|---|---|---|---|
| **PEG-N-PROXYL** | 15.93 G | - | - |
| **K16-PEG-N-PROXYL** | - | 15.33 G | 16.00 G |
| **K16-PEG-S-PROXYL** | - | 15.49 G | 16.02 G |

#### References


Hubbell WL, McConnell HM (1968) Spin-label studies of the excitable membranes of nerve and muscle. Proc Natl Acad Sci USA 61(1):12-16. doi:10.1073/pnas.61.1.12

Marsh D (2001) Polarity and permeation profiles in lipid membranes. Proc Natl Acad Sci USA 98(14):7777-7782. doi:10.1073/pnas.131023798
