## Supplementary figures and images for "Design of colipids with controlled lateral distance of head group moieties for targeting strategies in the field of lipid-based nanoparticles"

### Scheme 1 in high quality

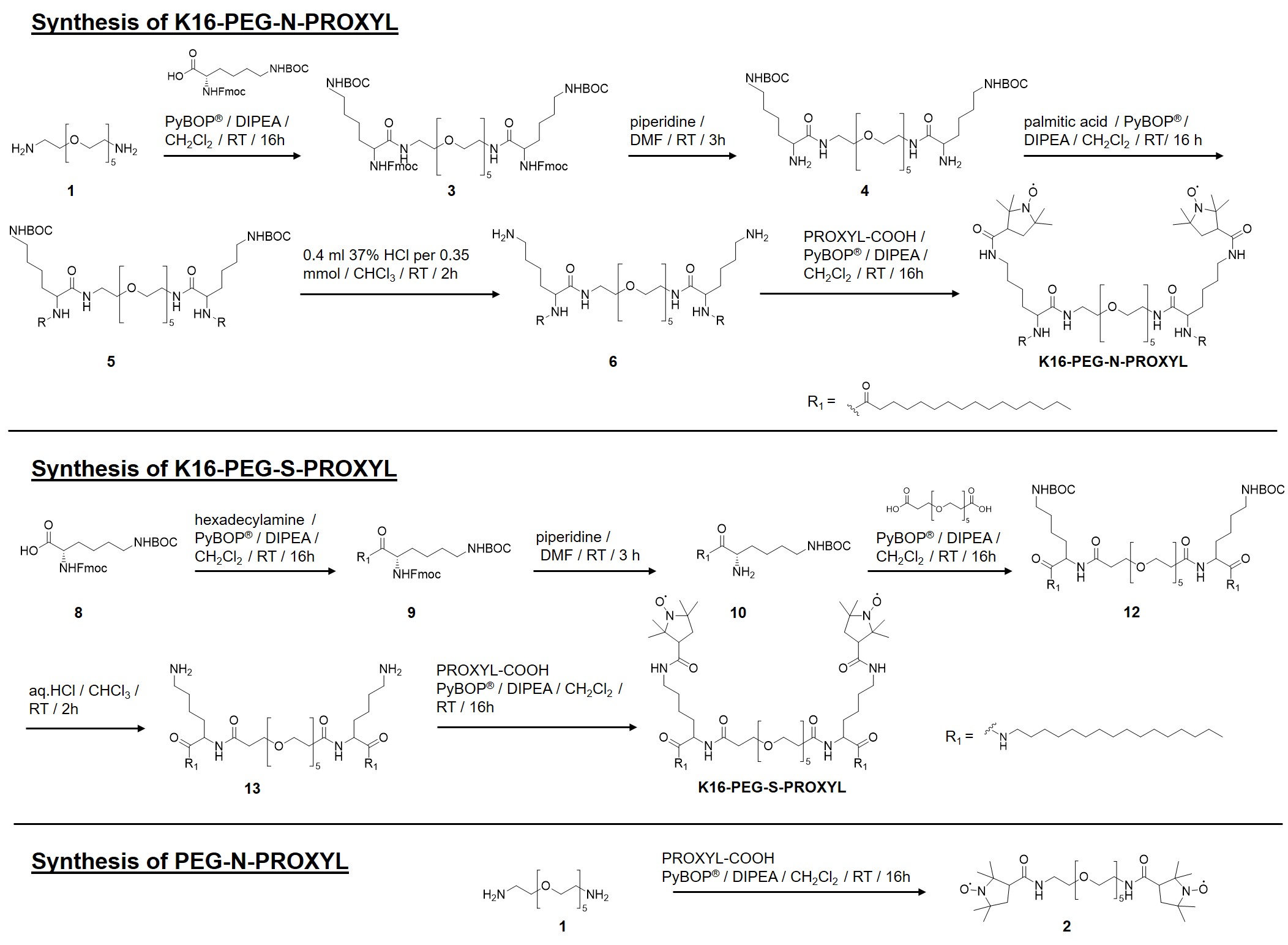
